# Insulin-like peptide secretion is mediated by peroxisome-Golgi interplay

**DOI:** 10.1101/2025.05.26.656179

**Authors:** Marie A. König, Nicole Kucharowski, Darla P. Dancourt Ramos, Hannah Soyka, Klaus Wunderling, Torsten R. Bülow, Mohamed H. Yaghmour, Christoph Thiele, Jan M. Ache, Lars Kuerschner, Margret H. Bülow

## Abstract

Insulin is a peptide hormone that is secreted in Golgi-derived dense-core vesicles from mammalian pancreatic beta-cells in response to nutrients. In *Drosophila melanogaster*, three insulin-like peptides are secreted as neuropeptides from the insulin-producing cells in the brain. Peroxisomes are lipid-metabolizing organelles that engage into various membrane contact sites with other organelles. Impaired peroxisomal metabolism has been associated with beta-cell apoptosis and impaired insulin secretion. How peroxisomes contribute to insulin and neuropeptide secretion is unknown. Here we demonstrate that peroxisomes interact with the Golgi apparatus in *Drosophila* insulin-producing cells. Secretion of insulin-like peptide 2 is cell-intrinsically impaired in mutants lacking the peroxisome assembly factor Pex19. Loss of peroxisomes shifts the profile of sphingolipids towards longer sphingoid bases and leads to accumulation of sphingolipids in the Golgi. We show that peroxisomes dynamically interact with the Golgi in insulin-producing cells and that Pex19 directly contributes to peroxisome-Golgi interaction via the fatty acyl-CoA reductase FAR2/waterproof in the peroxisomal membrane. We propose that this peroxisome-Pex19-Golgi axis is required to adjust Golgi membranes upon starvation by withdrawing lipids with longer side chains, thereby optimizing Golgi membrane flexibility for dense-core vesicle secretion upon refeeding.

## Introduction

Neuropeptide secretion is a fundamental process in neuronal communication, yet key aspects of its regulation at the subcellular level remain poorly understood. In *Drosophila melanogaster*, insulin-like peptides (Dilps) function not only as metabolic hormones but also as neuropeptides, given that insulin secretion occurs in specialized neurons rather than in a pancreas. The *Drosophila* genome encodes eight insulin-like peptides (Semaniuk et al., 2021), with both the insulin receptor (dInR) and Dilps displaying strong homology to their mammalian counterparts (Brogiolo et al., 2001). Among these, Dilp2, Dilp3, and Dilp5 are produced and secreted by insulin-producing cells (IPCs) in the pars intercerebralis, a neuroendocrine region of the central nervous system (J. Kim & Neufeld, 2015). Their secretion is tightly regulated: during starvation, Dilps accumulate in neuronal somata, while refeeding triggers their rapid release into the hemolymph, driving systemic metabolic adaptation (Barry & Thummel, 2016). Secretion of Dilps from IPCs is regulated by several cytokines and peptide hormones derived from the fat body, the adipose tissue and liver of the fly (Bai et al., 2012; Rajan & Perrimon, 2012; Sano et al., 2015). The cell-intrinsic mechanisms governing Dilp secretion from these neurons remain unresolved.

In mammalian pancreatic β-cells, the key subcellular structure in insulin secretion is the Golgi apparatus. In the Golgi, pro-insulin is packed into secretory vesicles (dense-core vesicles, DCV), where it is cleaved to yield the mature insulin (Ho et al., 2023). Inhibition of insulin secretion leads to changes in Golgi morphology (Iwamoto et al., 2023).

Peroxisomes, key regulator of lipid metabolism, are best known for their role in the β-oxidation of very-long-chain fatty acids (VLCFA) and the maintenance of cellular lipid homeostasis (Kumar et al., 2024). Loss of peroxisomes leads to profound metabolic imbalances, including lipid perturbations that contribute to mitochondrial dysfunction, decreased DCV abundance, and β-cell failure in the mouse pancreas (Baboota et al., 2019). In mammals, peroxisomal dysfunction is linked to insulin secretion defects, suggesting an evolutionarily conserved role in hormone regulation. Similarly, peroxisomes facilitate lipid remodeling in plants and fungi, underscoring their broader significance in cellular metabolism (Maruyama & Kitamoto, 2013; Reumann & Bartel, 2016). Despite their well-established metabolic functions, a direct role for peroxisomes in neuropeptide secretion remains unexplored.

In parallel, the Golgi-apparatus is a central hub for the post-translational modification, sorting, and secretion of DCVs. A defining feature of Golgi membranes is their dynamic lipid environment, where ceramide metabolism plays a crucial role in membrane integrity, vesicular trafficking, and protein sorting (Chitkara & Atilla-Gokcumen, 2025). Ceramide, synthesized in the endoplasmic reticulum (ER) and processed in the Golgi, governs membrane curvature and vesicle budding, in part through interactions with phosphatidylinositol-4-phosphate (PI4P) (van Galen et al., 2014). Perturbations in ceramide homeostasis disrupt neuropeptide secretion, highlighting its importance in regulated exocytosis (Magnan & Le Stunff, 2021). Beyond sphingolipid metabolism, the biophysical properties of membranes, which are influenced by the chain length of fatty acid (FA) incorporated in its lipids, shape the secretion process. Shorter-chain FAs reduce van der Waals interactions, thereby increasing membrane fluidity, while longer chains promote tighter packing and rigidity (Bao et al., 2021). The Golgi apparatus exhibits a gradient of increasing FA chain length from cis- to trans-Golgi, correlating with functional compartmentalization and vesicle maturation (Agliarulo & Parashuraman, 2022).

A direct contribution of peroxisomes to insulin or neuropeptide secretion has not yet been established. Here, we demonstrate that loss of the peroxisome biogenesis factor Pex19 leads to impaired Dilp secretion from IPCs following a feeding stimulus. This impairment is driven by the incorporation of longer fatty acids into sphingolipids, altering Golgi membrane composition and vesicle dynamics. Our study demonstrates that Pex19 plays a crucial role in mediating Golgi-peroxisome contacts. This interaction has been described in yeast, where peroxisomal association with the Golgi increases under amino acid starvation (Castro et al., 2022), and in human cells (Lee et al., 2024), but a function for this interaction has not been established previously. Here we provide evidence for functional peroxisome-Golgi interactions in a metazoan and thereby present a novel mechanism for a major biological process in a cell, namely secretion. We show that these contacts are nutrient-responsive in *Drosophila* IPCs, being abundant during starvation but significantly reduced upon refeeding. Our findings suggest that peroxisomes interact with the Golgi via Pex19, the peroxisomal fatty acyl-CoA reductase FAR2/wat, and its interactors in the Golgi membrane, and with lipidomics analysis of DCVs we show that this interaction is required to metabolize VLCFAs from Golgi lipids, thereby altering Golgi membrane fluidity and allowing neuropeptide secretion upon refeeding.

Together, our data reveal a previously unrecognized role for peroxisomes in neuropeptide secretion, providing a mechanistic link between peroxisomal lipid metabolism and DCV release. By identifying peroxisome-Golgi interactions as key regulatory axis, we establish peroxisomes as active players in neuropeptide secretion and metabolic signaling.

## Results

### Dilp2 secretion from larval IPCs is impaired in Pex19 mutants

We showed previously that *Drosophila* mutants for the peroxisome assembly factor Peroxin 19 (Pex19) reflect hallmarks of human peroxisomal biogenesis disorders (PBDs) including accumulation of VLCFA, mitochondrial damage and neurodegeneration (Bülow et al., 2018; Sellin et al., 2018). We characterized disturbed lipid metabolism in Pex19-deficient flies and human cells and identified a shortage in medium-and long-chain fatty acids (MCFA/LCFA) rather than VLCFA as a mortality-driving factor of peroxisome dysfunction. Pex19 mutants showed several indications of starvation, such as reduction of fat body tissue and transcriptional upregulation of the starvation markers Hnf4 and Lipase 3 (Bülow et al., 2018; Hänschke et al., 2022). To gain more insight into the metabolic perturbances elicited by peroxisome loss, we analyzed insulin secretion. Insulin-like peptides 2, 3 and 5 are produced in a set of peptidergic neurons in the pars intercerebralis of the *Drosophila* brain (Brogiolo et al., 2001), from where they are secreted in a nutrient-dependent manner and transferred to other neuron groups (Bader et al., 2013; Sarraf-Zadeh et al., 2013) or into the hemolymph to target peripheral tissues in both larvae and adults (Barry & Thummel, 2016; J. Kim & Neufeld, 2015) (Figure 1A). We used white mutants (w^1118^) as control and pex19 mutant larvae (in the w^1118^ genetic background) that were synchronized to the same developmental stage (early 3^rd^ instar) and starved them for two hours on PBS-soaked filter paper. We found that under this condition, Dilp2 is efficiently retained in the somata of the IPCs in both wildtype and pex19 mutants. Upon one hour of refeeding with 10 % sucrose and yeast paste, IPCs in wildtypes cleared their somata from Dilp2, suggesting efficient secretion. In pex19 mutants, Dilp2 remained at high levels (Figure 1B, D). We concluded that pex19 mutants cannot secrete Dilp2 from IPCs upon a nutrient stimulus. Similarly, the secretion of Dilp3 is impaired (Supplemental Figure 1A, B).

**Figure 1:**
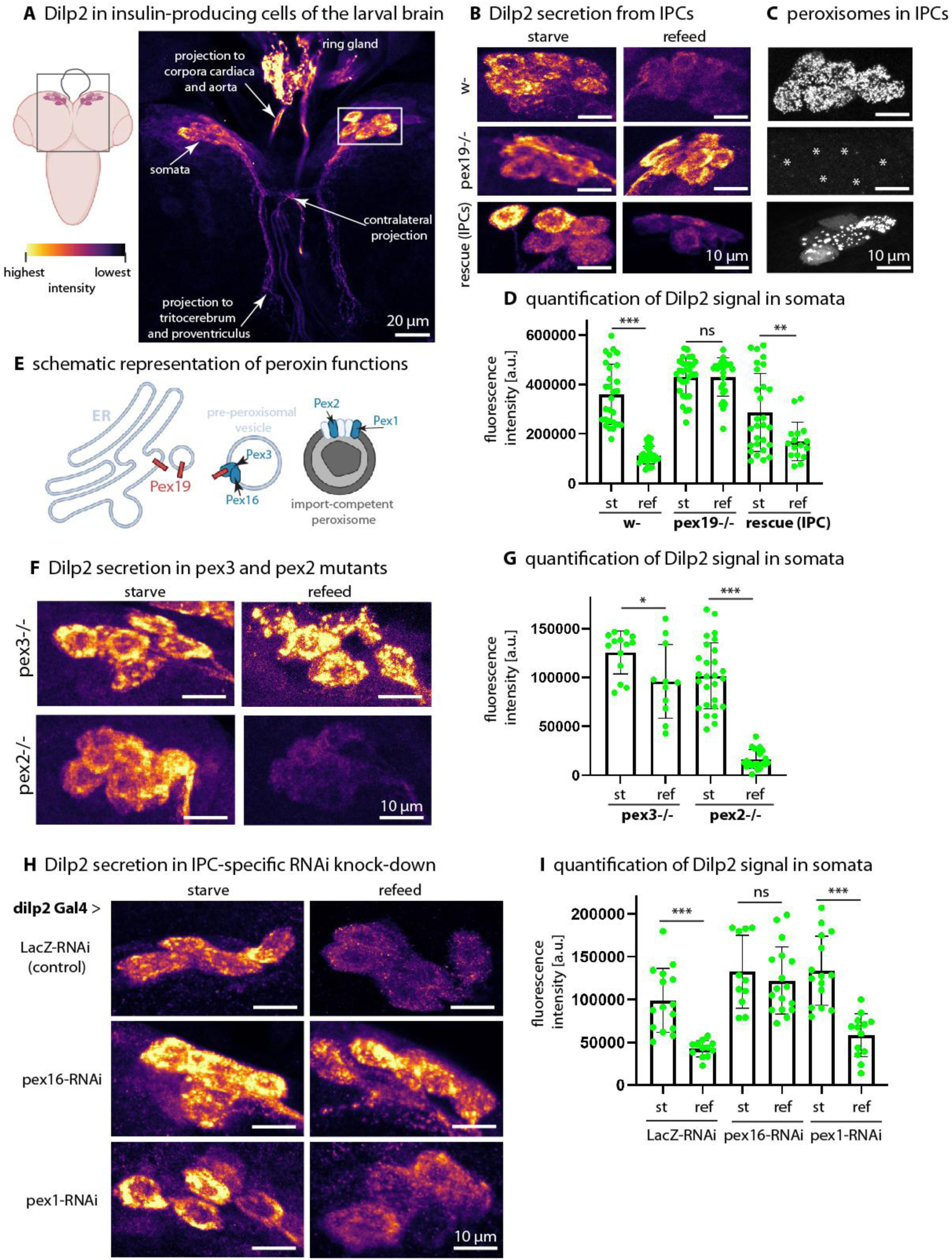
(A) Localization of insulin-producing cells (IPCs) and Dilp2 distribution in the *Drosophila* larval brain. Insulin-like peptides Dilp2, Dilp3, and Dilp5 are secreted from a set of peptidergic neurons in the pars intercerebralis, a neuroendocrine center functionally analogous to the mammalian hypothalamus. In larvae, IPCs extend projections to the corpora cardiaca and the tritocerebrum (subesophageal zone), regions involved in metabolic regulation and neuronal communication. Heatmap indicates Dilp2 fluorescence intensity (highest to lowest). (B) Dilp2 secretion from IPCs under starvation and refeeding conditions in wild-type (w⁻), pex19 mutant (pex19⁻/⁻, pex19^ΔF7/ΔB2^), and IPC-specific rescue (w-; pex19⁻/⁻, UAS Pex19; dilp2,3 Gal4) larvae. (C) Peroxisomes in IPCs, visualized by GFP-SKL. Asterisks indicate cells lacking peroxisomes. (D) Quantification of Dilp2 fluorescence intensity in IPC somata across genotypes and nutritional states. (E) Schematic representation of peroxin functions, illustrating the roles of Pex19, Pex3, and Pex16 in peroxisome biogenesis, while Pex1 and Pex2 function in matrix protein import to maintain peroxisomal metabolic activity. (F) Dilp2 secretion in pex3 (pex3^2^/Df(32)6262) and pex2 (pex2^HP35039^/pex2^f01899^) mutants under starvation and refeeding conditions. (G) Quantification of Dilp2 fluorescence intensity in IPC somata of pex3 and pex2 mutants across conditions. (H) Dilp2 secretion in IPC-specific (*dilp2*-Gal4) RNAi knockdown larvae, comparing control (LacZ-RNAi), pex16-RNAi, and pex1-RNAi under starvation and refeeding conditions. (I) Quantification of Dilp2 fluorescence intensity in IPC somata of RNAi knockdown larvae. Scale bars as indicated. Asterisks represent * p < 0.05, **p < 0.01, *** p < 0.001 and ns (not significant).

### Dilp2 secretion is impaired in pex19 mutants in a cell-autonomous manner

We showed that in mutants for the peroxisome assembly factor Pex19, Dilps are retained in the soma upon starvation, but are not released upon a refeeding stimulus. Secretion of Dilps is stimulated by systemic signals, such as factors that derive from the fat body, an endocrine organ with functional homology to the mammalian liver and adipose tissue. The fat body secretes e.g. Dilp6 and Unpaired-2 (upd2), a cytokine, which reach the IPCs via the hemolymph to regulate the release of Dilps (Bai et al., 2012; Rajan & Perrimon, 2012). To find out if systemic loss-of-function (LOF) of Pex19 abrogates endocrine signaling to the IPCs and thereby prevents Dilp release, we reconstituted Pex19 specifically in IPCs (Figure 1B, C, lower panel). Mature, import-competent peroxisomes can be visualized with a peroxisome-targeted GFP (GFP-SKL). When expressed in wildtypic IPCs, peroxisomes are visible as small, soma-restricted dots (Figure 2C). This signal is absent in the pex19 mutant. Upon reconstitution, peroxisomes are visible as discrete dots in IPCs, and somata clearing of Dilps upon a nutrient stimulus is restored. This demonstrates that the defect in Dilp2 secretion in pex19 mutants is a cell-intrinsic process. Strikingly, this IPC-specific reconstitution partially rescues their survival, yielding ∼40 % adult flies (Supplemental Figure S1C). Systemic reconstitution of Pex19 in hemocytes (*hml*-Gal4) also rescues their survival, yielding ∼60 % adult flies, but does not rescue Dilp2 secretion (Supplemental Figure S1D, E).

**Figure 2:**
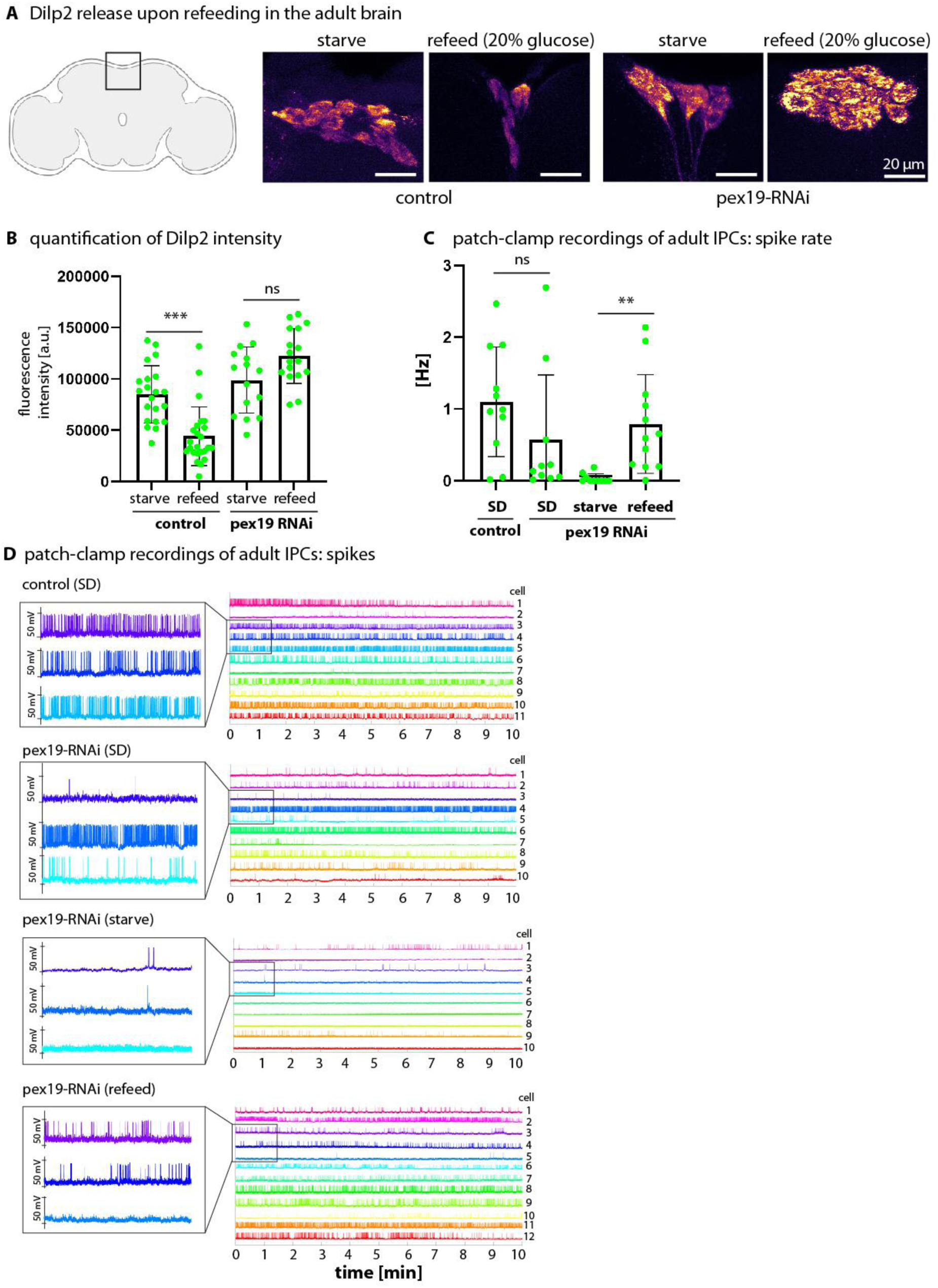
(A) Representative images of adult fly brains showing Dilp2 accumulation and release in insulin-producing cells (IPCs) under different nutritional conditions (starvation and refeeding) in control (dilp2 Gal4/ UAS-mCD8-GFP) and Pex19 IPC-specific knock-down flies (dilp2 Gal4/ UAS-mCD8-GFP; UAS-Pex19-RNAi). (B) Quantification of Dilp2 fluorescence intensity in IPCs across conditions. (C) Spike rate of IPCs activity in patch-clamp recordings (spike frequency in Hz) in control and Pex19-RNAi flies under *ad libitum* fed conditions (standard diet, SD), 24 h starved and 4 h refed on 20 % glucose. (D) Patch-clamp recordings of spontaneous IPC activity in control and Pex19-RNAi flies. Left panels show spike amplitudes in representative trials, while right panels overlay > ten trials per condition. Scale bars as indicated. Asterisks represent * p < 0.05, ***p < 0.001, ***p < 0.0001 and ns (not significant).

### Dilp2 secretion upon refeeding is impaired in pex19 and pex3, but not in pex2 mutants

Peroxisome assembly is mediated by peroxin proteins of three categories: Pex3, Pex16 and Pex19 are required for the import of membrane proteins already in early stages of peroxisome *de novo* biogenesis. Thus, depletion of these factors leads to the absence of peroxisomal membranes. Peroxins such as Pex1, Pex2, Pex6 etc. are required for the import of matrix proteins that are recognized by their peroxisomal targeting sequence (Figure 1E). Theoretically, upon depletion of these factors and in the presence of Pex3, Pex16 and Pex19, remnants of peroxisomal membranes, referred to as “ghosts”, can be present. We asked if Dilp2 secretion would also be affected in other peroxin mutants. In mutants for the Pex19 receptor, Pex3, Dilp2 secretion upon refeeding is blocked, similar to what we observed in pex19 mutants (Figure 1F, G). In mutants for a matrix protein import factor, Pex2, we did not observe this phenotype: upon refeeding, Dilp2 is efficiently cleared from IPCs of pex2 mutants (Figure 1F, G).

### Membrane peroxins, but not matrix peroxins, contribute to Dilp2 secretion

We asked if RNAi-mediated knock-down of peroxin proteins in IPCs would be sufficient to block Dilp2 release. In an RNAi control (LacZ-RNAi), Dilp2 is efficiently released upon refeeding (Figure 1H, I). When Pex16, a protein that like Pex3 and Pex19 is required for membrane protein import, is knocked down, Dilp2 secretion is blocked similar to what we observed in pex19 mutants (Figure 1H, I). This supports our finding that peroxisomal membrane assembly factors are required for Dilp2 secretion. By contrast, knock-down of Pex1, a protein involved in matrix protein import, was not sufficient to block Dilp2 secretion upon refeeding (Figure 1H, I). In sum, our data suggest that peroxisomal membrane assembly factors, but not factors that mediate peroxisome import, are required for the secretion of insulin-like peptides from IPCs.

### Ablation of Pex19 in IPCs of adults impairs Dilp2 secretion but not electrophysiological activity

Next we asked if Dilp2 secretion is impaired under the loss of Pex19 due to physiological inactivity. We used patch-clamp recording of the electric currents in these neurons and determined the activity as the spike frequency (Hz) in adult animals (Liessem et al., 2023). First, we confirmed that knock-down of Pex19 is sufficient to block Dilp2 release upon refeeding in brains of adult flies. We found that in control adults, Dilp2 accumulates in IPCs upon 24 h of starvation and is efficiently released upon 4 h refeeding with 20 % glucose. IPC-restricted RNAi-mediated knock-down blocks Dilp2 release upon refeeding (Figure 2A, B). We then performed patch-clamp recordings of IPCs using the same genotypes. *Ad libitum* fed control flies (expressing mCD8-GFP in IPCs) show high neuronal activity, represented as the spike rate recorded from more than ten cells (Figure 2C, D). IPC-restricted knock-down of Pex19 in *ad libitum* fed flies show slightly reduced activity. As shown in (Bisen et al., 2025), IPCs shut down their electrophysiological activity upon starvation and gradually reactivate the cells upon refeeding with glucose. Pex19 knock-down leads to almost complete inactivity of IPCs (Figure 2C, D, 3^rd^ panel). Upon refeeding with 20 % glucose, the electrophysiological activity is restored (Figure 2C, D, lower panel). We thus conclude that the loss of Pex19 does not impair the electrophysical activity of IPCs. Surprisingly, this suggests that Dilp secretion and neuronal activity are uncoupled under Pex19 loss-of-function in IPCs.

### Medium- and long-chain fatty acid decrease affects sphingolipids

A major hallmark of peroxisome loss is the accumulation of VLCFA, substrates of the peroxisomal β-oxidation machinery. We showed previously that the FA profile of pex19 mutants exhibits a marked accumulation of VLCFA and a decrease in M- and LCFA, in particular of C14 FAs (Bülow et al., 2018, Sellin et al., 2018). Quantification of FA methylesters revealed that M- and LCFA are an order of magnitude more abundant than VLCFA, leading to an overall shift in the average FA chain towards longer chains in pex19 mutants. Here we measured the lipidome of control and pex19 mutant larvae. Expectedly, we found that the ether phospholipid PE-O (plasmalogen), a peroxisomal product, is reduced (Figure 3A’). We observed a strong increase in phosphatidylinositol (PI) from ∼ 500 pmol/mg wet weight in controls to ∼ 1300 pmol/mg in pex19 mutants (Figure 3A’’). Note that control (w-) values of PI and PE-O were already shown in (Hänschke et al., 2022). The phospholipids phosphatidylethanolamine (PE), phosphatidylglycerol (PG) and phosphatidylserine (PS) were not significantly altered, nor were cholesterylester and the neutral lipid diglyceride (DG, Supplemental Figure S2A). Triglyceride (TG) was elevated in pex19 mutants (Supplemental Figure S2A). Among the sphingolipids, ceramide (Cer) was unchanged, but ceramide phosphatidylethanolamine (CerPE) and sphingomyelin (SM) were reduced and Hexosylceramide (HexCer) was elevated in pex19 mutants (Figure 3A’’’). We then analysed the distribution of lipid species in the individual classes and found that the shift towards longer carbon chains is not equally represented in all lipid classes, but is pronounced in sphingolipids where it is accompanied by an increase in the length of the spingoid base. In ceramide (Figure 3B’), HexCer (Figure 3B’’), SM (Figure 3B’’’) and CerPE (Figure 3B’’’’), we find a marked shift from lipid species with a C14 base in control larvae to a C16 base in pex19 mutants, which parallels the overall shift towards longer FA chains that we described earlier. A similar phenomenon has been described in *Drosophila* mutants for Pex16 and Pex2 (Wangler et al., 2024). We have shown earlier that dietary administration of MCFA (by addition of 5% coconut oil to the diet) rescues the lethality of the pex19 mutation. Here we tested Dilp2 secretion in pex19 mutants that were fed with MCFA prior from the first larval stage to starvation onset and found that it rescues also Dilp2 secretion upon refeeding (Figure 3C, D). This suggests that the shift in sphingolipids towards longer chains is connected to the defect in Dilp2 secretion.

**Figure 3:**
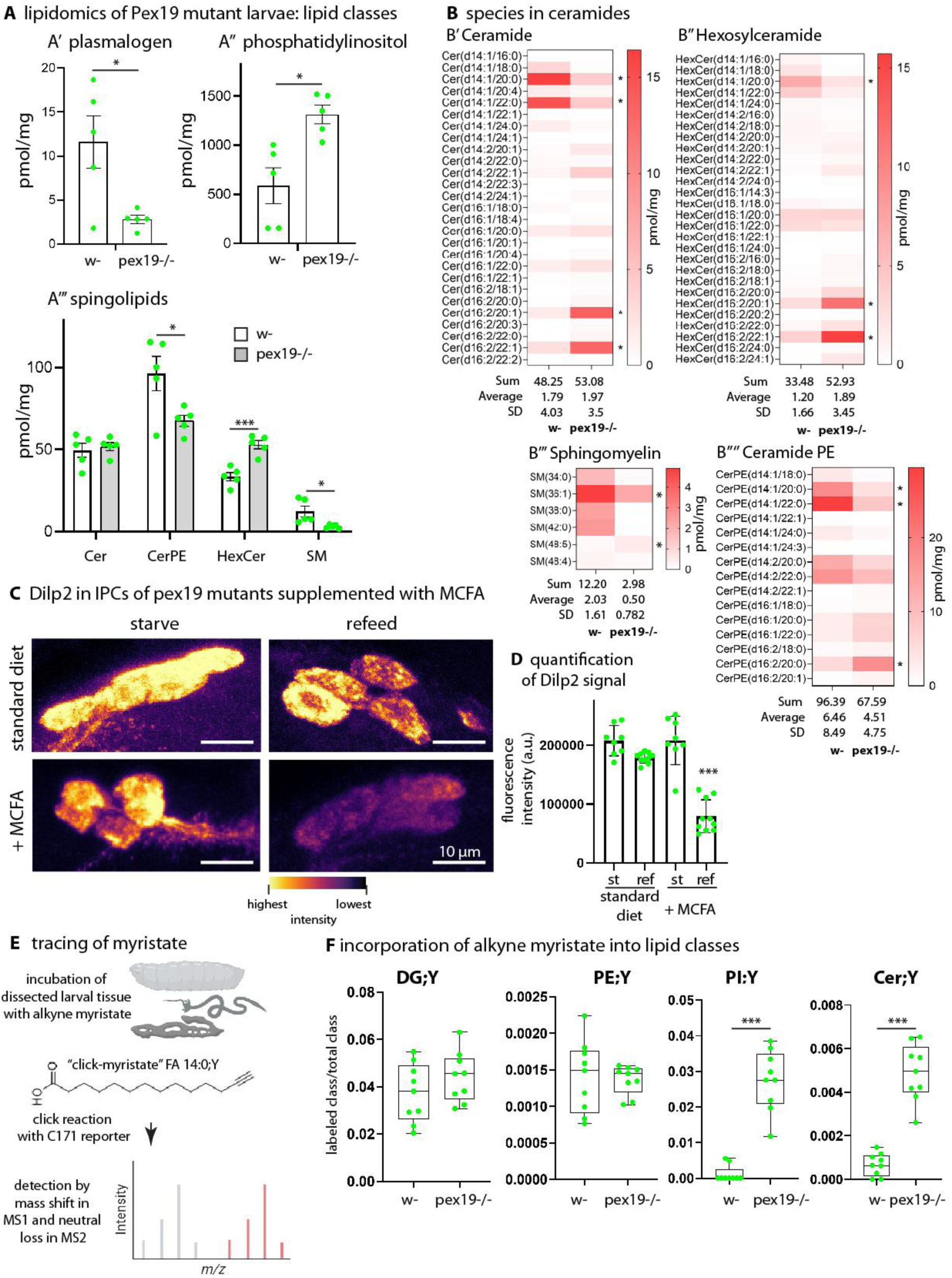
(A) Lipidomics of w- and pex19 mutant larvae separated by different lipid classes reveals (A’) a characteristic loss of plasmalogens (PE-O) in pex19 mutants due to peroxisomal loss, (A’’) a significant increase in phosphatidylinositol (PI) levels upon pex19 mutation, and (A’’’) notable alterations in sphingolipid profiles. (B) Analysis of ceramide species shows sphingolipid side chain length distribution, with (B’) ceramides (Cer), (B’’) hexosylceramide (HexCer), and (B’’’) ceramide phosphoethanolamine (CerPE) in control versus pex19 mutants. The data reveal a shift from shorter FA chains and sphingoid bases (C14) in control larvae to longer FA chains and bases (C16) in pex19 mutants. (C) Dilp2 secretion in IPCs of pex19 mutants supplemented with medium-chain fatty acids (MCFAs) shows rescue of the secretion defect induced by pex19 mutation, (D) with corresponding quantification of fluorescent intensity. (E) Schematic representation of alkyne-tagged myristate (FA14:0;Y) tracing in dissected larval tissue, followed by mass spectrometry-based detection of incorporation through mass shift analysis. (F) Incorporation of alkyne myristate into lipid classes reveals no significant differences in diacylglycerol (DG) or phosphatidylethanolamine (PE) between control and pex19 mutant larvae, but highly significant incorporation into PI and Cer in pex19 mutants. Scale bars as indicated. Asterisks represent * p < 0.05, ***p < 0.001 and ***p < 0.0001.

### A myristate tracer shows a requirement for C14 chains in phosphatidylinositol and ceramide

To understand which lipid classes are affected by the decrease in MCFA, we performed a lipid tracing experiment (Figure 3E). We incubated permeabilized larval tissue with a traceable analogue of myristic acid, the alkyne FA 14:0;Y, and followed its incorporation into the lipidome after a click reaction (Thiele et al., 2019). Control and pex19 mutant samples showed similar incorporation of the tracer into lipids such as labeled DG;Y or PE;Y, but increased tracer incorporation into labeled PI;Y (phosphatidylinositol), the precursor of the signalling lipids phosphoinositides (Figure 3F), in pex19 mutants. However, we did not observe a shift towards longer average side chains in PI (Supplemental Figure S2B). Strikingly, also incorporation of alkyne myristate into labeled ceramide (Cer;Y) is increased in pex19 mutants, suggesting that phosphatidylinositol and ceramide are primarily affected by the decrease in C14. The intriguing increase in (unlabeled) PI could be linked to altered levels of phosphoinositides that are important for insulin signaling. Following this hypothesis, we overexpressed PI3K and PTEN, enzymes that catalyze the conversion of PIP2 to PIP3 and back to PIP2, respectively, but found no change in the nutrient-dependent secretion of Dilp2 (Supplemental Figure S2C). We overexpressed phospholipase C δ (PLC-δ) with a GFP coupled to its plecstrin homology domain (PLC-δ-PH-GFP). PLC-δ catalyzes the conversion of PIP2 to diacylglycerol and inositoltrisphosphate. Also here, we did not find an impairment of Dilp2 release (Supplemental Figure S2D). We thus continued to investigate the role of ceramide in impaired Dilp2 secretion of pex19 mutants.

### Golgi sphingolipids accumulate in PEX19 mutant HeLa cells and Drosophila tissue

Insulin-like peptides are packed into dense-core vesicles (DCVs) at the trans-Golgi network. Golgi membranes are rich in ceramides; in fact, BODIPY-ceramide is a probe that serves as a Golgi marker (Bülow & Broichhagen, 2025). We stained control and PEX19 knock-out HeLa cells with BODIPY-Cer (Schrul & Kopito, 2016) and labeled peroxisomes with a PMP70 antibody to show that peroxisomes are absent from these cells (Figure 4A, B). BODIPY-Cer heavily accumulates in PEX19-deficient cells. We used another ceramide-based Golgi probe, NBD-C6-ceramide, and found that its Golgi signal increases in PEX19-deficient HeLa cells (Figure 4C, D). Next, we tried BODIPY-Cer in larval tissue of *Drosophila* (Figure 4E). In fat body and gut tissue of control animals, we found that the probe labels the plasma membrane rather than subcellular structures. In pex19 mutants, the signal is drastically increased, suggesting that also here, ceramides accumulate. In sum, we found that ceramide-based Golgi probes heavily accumulate in pex19 mutant gut and fat body tissue as well as in HeLa cells carrying a PEX19 mutation. However, we do not observe a general accumulation of ceramide in the lipidome of whole larvae. We thus conclude that ceramides accumulate in the Golgi apparatus but are not distributed to other membranes in pex19 mutants, leaving overall ceramide levels unaffected.

**Figure 4:**
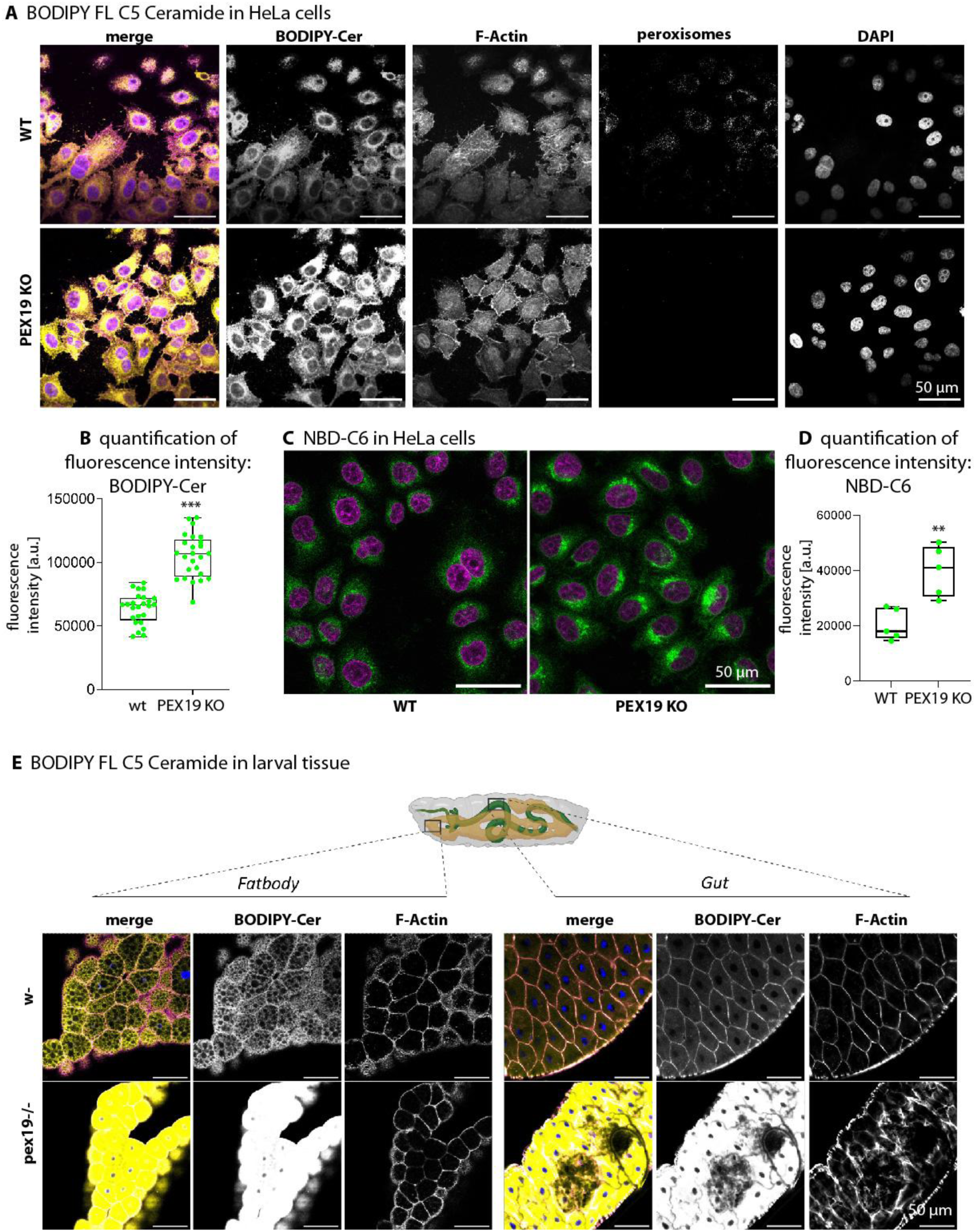
(A) Fluorescent imaging of BODIPY FL C5-Ceramide (BODIPY-Cer) in HeLa cells reveals increased ceramide accumulation in PEX19 knockout (KO) cells compared to wild-type (WT), with preserved nuclear and actin architecture and missing or intact peroxisomal structures, respectively. F-Actin is labeled with phalloidin-Alexa555 and peroxisomes with an antibody against peroxisomal membrane protein 70 (PMP70). (B) Quantification of BP-C5 fluorescence intensity confirms a highly significant increase in PEX19 KO cells relative to WT. (C) Imaging with NDB-C6, a Golgi-targeting ceramide analog, shows enhanced Golgi-associated ceramide accumulation in PEX19 KO cells, with (D) corresponding quantification indicating a significant elevation in fluorescent signal. (E) Schematic representation of larval tissue and fluorescent images of dissected larval gut and fat body tissue from pex19 mutant and control larvae demonstrate striking ceramide accumulation in mutant tissues, while actin staining indicates intact cellular morphology. Scale bars as indicated. Asterisks represent **p < 0.01, ***p < 0.001.

### Peroxisomes interact with the Golgi apparatus in a nutrient-dependent manner

Next we asked if peroxisomes directly interact with the Golgi apparatus in IPCs, an organelle interaction that has not been shown in metazoans yet. We used different combinations of markers for peroxisomes and Golgi with fluorescent tags: GFP-SKL, a GFP with a minimal peroxisomal targeting signal and Golgi-RFP (carrying a galactosyl-transferase tagged with RFP at the N-terminus). We created a peroxisome-mcherry (px-mcherry) line carrying an mcherry construct with a C-terminal peroxisomal targeting sequence (PTS1) from the peroxisomal enzyme CRAT (carnitine-O-acetyltransferase (Faust et al., 2012)). We used it in combination with Grasp65-GFP to label the Golgi apparatus. Using Airyscan super-resolution microscopy, we could observe that under starvation, many peroxisomes are in close proximity with the Golgi and show colocalization, suggesting that they engage in contacts with the Golgi (Figure 5A, B). We also tested a different combination of markers, GFP-SKL (SKL as the minimal peroxisomal targeting sequence) and Golgi-RFP (galactosyltransferase-RFP) and analyzed them by super-resolution microscopy (Figure 5A, B). We then established a pipeline to quantify peroxisome-Golgi proximity with the software Imaris. Using the combination GFP-SKL, Golgi-RFP, we used the Imaris pipeline to show peroxisome-Golgi interaction in the “masked signal” of super-resolution images, which illustrate decreased interaction upon refeeding (Figure 5C). Upon quantification, we observed the same trends (Figure 5D) and a significant decrease in the ratio of peroxisomes colocalizing with Golgi membranes upon refeeding (Figure 5E). We then used the Imaris pipeline to perform a 3D reconstruction of Golgi membranes and contact-dependent color-coding of peroxisomes of confocal z-stacks. These show peroxisomes that are in contact with the Golgi in magenta, and peroxisomes without contact in green. Under refeeding, the number of peroxisomes that are in contact with Golgi membranes decreases (Figure 5F and Supplemental Figure 3C). Thus, peroxisome-Golgi contacts in IPCs are dynamic and depend on the feeding status of the animal.

**Figure 5:**
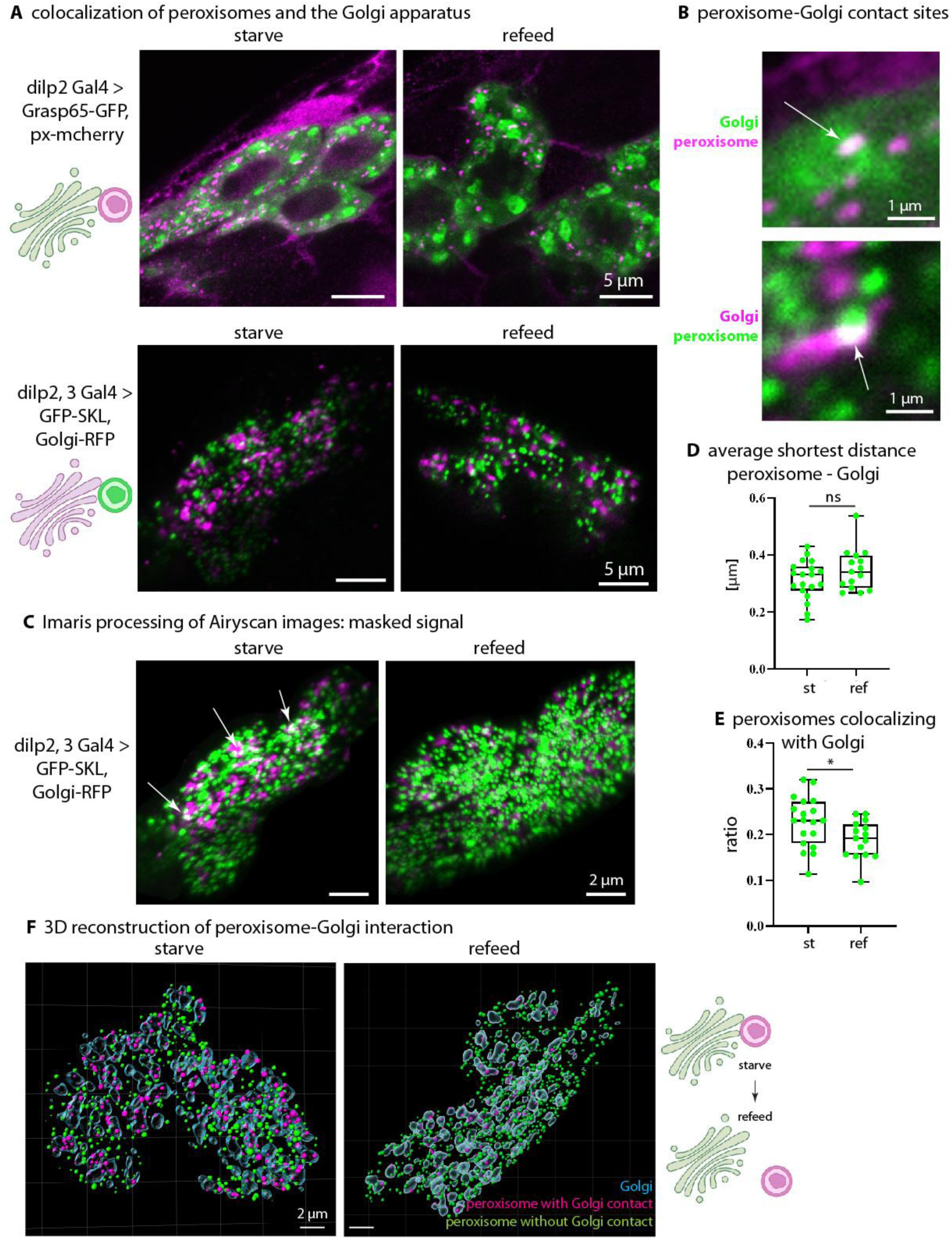
(A) Colocalization of peroxisomes and the Golgi apparatus in insulin-producing cells (IPCs). Single confocal planes of Airyscan super-resolution images from dilp2 Gal4, UAS Grasp65-GFP; UAS peroxi-mcherry and +; dilp2,3 Gal4, GFP-SKL/Golgi-RFP larvae under starvation and refeeding. (B) Peroxisome-Golgi colocalization sites. Close-ups of single confocal planes of Airyscan super-resolution images. (C) masked signal of Imaris 3D rendering images from Airyscan super-resolution z-stacks, genotype +; dilp2,3 Gal4, GFP-SKL/Golgi-RFP. Arrows point to peroxisome-Golgi colocalization. (D) Quantification of the average shortest distance between peroxisomes and Golgi in IPC clusters, comparing starved and refed states, genotype +; dilp2,3 Gal4, GFP-SKL/Golgi-RFP. (E) Ratio of peroxisomes colocalizing with the Golgi (peroxisomes with shortest distance ≤ 0 μm), genotype +; dilp2,3 Gal4, GFP-SKL/Golgi-RFP. Each data point represents the average of one IPC cluster. (F) 3D reconstruction of confocal Z-stacks. Surface rendering models generated using Imaris, displaying peroxisomes (colored dots) in relation to the Golgi (blue surface) in starved and refed states. Peroxisomes are color-coded based on their shortest distance to the Golgi, purple (with Golgi contact) and green (without Golgi contact). Scale bars as indicated. Asterisks represent * p < 0.05, ns: not significant.

### Dense-core vesicle lipidomics reveal loss of lipid dynamics upon refeeding in pex19 mutants

The trans-Golgi network secretes neuropeptides (such as insulin-like peptides) and peptide hormones (such as insulin) in DCVs. In order to test if DCVs would react to refeeding by altering their properties, and if this response would be missing in pex19 mutants, we adapted a protocol (Birinci et al., 2020) to enrich DCVs from larval extracts using a sucrose gradient. To show that the protocol can enrich DCVs from *Drosophila* larvae, we tested if Dilp2 accumulates in the final pellet of the protocol. We compared larval extracts (input) and the DCV-containing pellet and found that Dilp2 is enriched. We used the adapted protocol to enrich DCV from control and pex19 mutant larvae from starved and refed conditions, isolated lipids and subjected them to lipidomics analysis. We found that DCV lipids from controls exhibited a nutrient-dependent dynamic: upon refeeding, the lipid content (normalized to protein content) decreased in PI, sphingolipids and also on the level of total lipids (Figure 6A, Supplemental Figure S4A). Strikingly, ceramides accumulate in enriched DCVs of pex19 mutants, supporting our hypothesis that Golgi-derived lipids accumulate but are not distributed to other membranes in pex19 mutants (Figure 6B, compare also Figure 4E). Many ceramide species with long or very long fatty acid side chains are reduced in controls, leading also to an overall reduction in ceramide upon refeeding. These species are at high levels in starved pex19 mutants and increase even further upon refeeding. In human cells, the Golgi matrix protein GM130 (encoded by *GOLGA2*) is predicted as a direct interactor of PEX19 (Huttlin et al., 2021). DCVs of pex19 mutants showed an accumulation of the Golgi matrix protein GM130 (Figure 6D, Supplemental Figure S4B).

**Figure 6:**
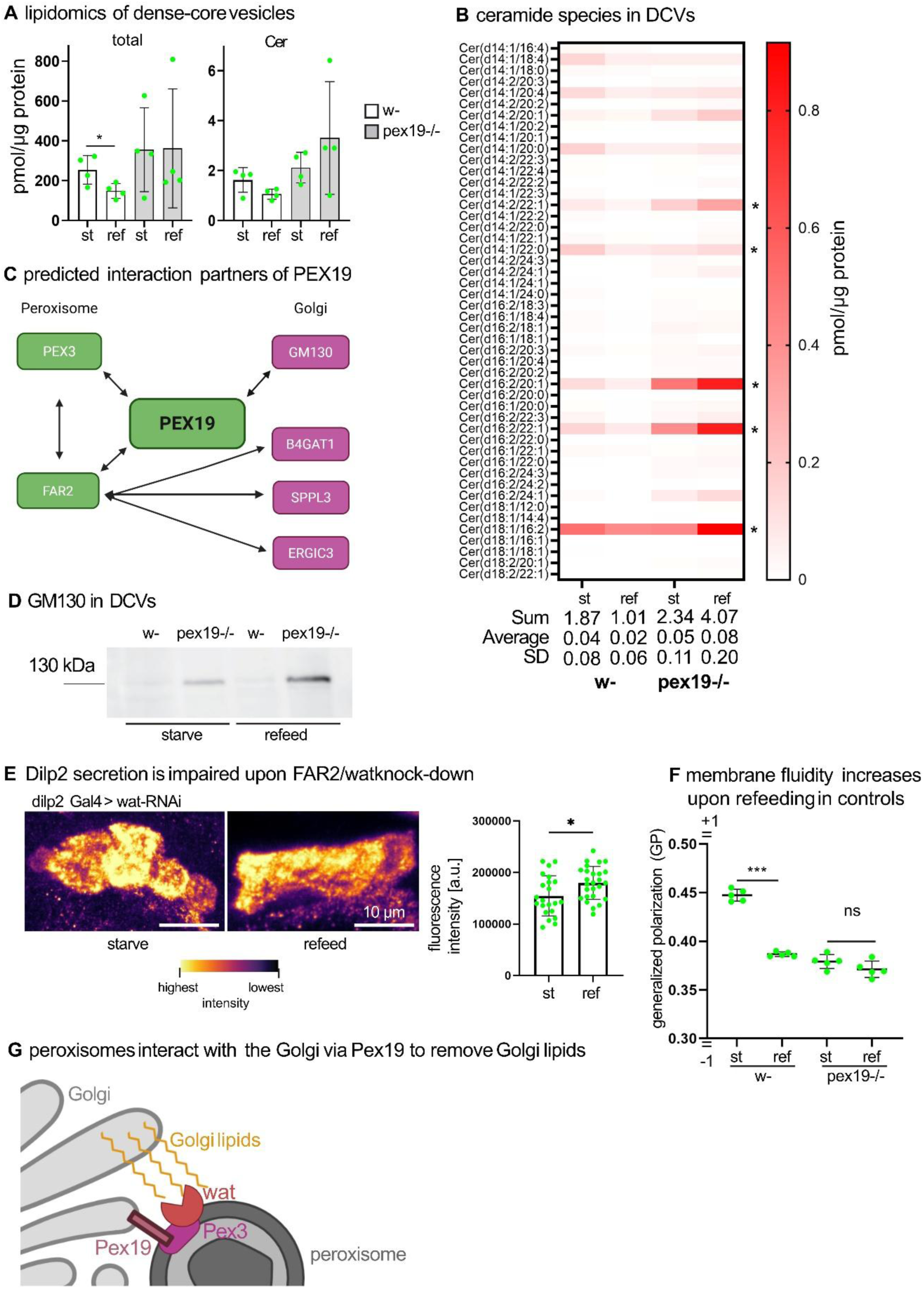
(A) Lipidomics analysis of enriched DCVs from control and pex19 mutants upon starvation and refeeding. Lipid classes as indicated and normalized to protein content. (B) Distribution of ceramide species of enriched DCVs from control and pex19 mutants upon starvation and refeeding. Asterisks mark species with long or very long fatty acids that decrease upon refeeding in controls but remain at elevated levels in pex19 mutants. (C) Schematic representation of PEX19 interaction partners in human cells. (D) Western Blot of GM130: the Golgi matrix protein GM130 accumulates in pex19 mutant DCVs. (E) RNAi-mediated knock-down of FAR2/waterproof impairs Dilp2 release upon refeeding. (F) Generalized polarization of enriched DCVs determined by the fluorescence of Laurdan. (G) Schematic representation of functional peroxisome-Golgi interaction. Scale bars as indicated. Asterisks represent * p < 0.05, ** p < 0.01, ***p < 0.001, ns: not significant.

This further supports our hypothesis that in the absence of Pex19, DCV sorting and secretion is impaired.

### The peroxisomal membrane enzyme fatty acyl-CoA reductase FAR2/waterproof is predicted to interact with Pex19 and Golgi membrane proteins and affects Dilp2 secretion

We showed earlier in this study that the block of Dilp2 secretion upon refeeding is restricted to peroxins that play a role in membrane protein import. In mutants for the matrix protein import factor Pex2 or upon IPC-specific knock-down of Pex1, Dilp2 secretion is not affected. We thus assumed that Pex19 had a specific role in regulating peroxisome-Golgi interaction and thereby enable Dilp2 secretion upon refeeding. We used bioGRID (https://thebiogrid.org/) to search for potential Pex19 interactors that would link peroxisome and Golgi membranes. Apart from the interaction with GM130, we identified the peroxisomal fatty acyl-CoA reductase FAR2 (or wat (waterproof) in *Drosophila* (Jaspers et al., 2014)) as a candidate. FAR2 catalyzes the reduction of fatty acyl-CoAs to fatty alcohols, thereby contributing to the metabolism of VLCFA (Honsho et al., 2013). FAR2 also interacts with several Golgi proteins (Figure 6C) (Huttlin et al., 2015, 2021; Yang et al., 2023). We knocked-down wat in IPCs by RNAi and found that release of Dilp2 upon refeeding was blocked (Figure 6E). We hypothesized that the interaction between peroxisomes and Golgi upon starvation would help in the removal of lipids, such as sphingolipids, and specifically of longer chain fatty acids from Golgi membranes, thereby facilitating DCV secretion upon refeeding. This is also supported by the accumulation of Golgi markers in pex19-deficient cells and tissue. To test whether lipid removal from DCVs upon refeeding would affect membrane fluidity, we performed a Laurdan assay on DCV-enriched samples. Laurdan (6-Dodecanoyl-2-dimethylaminonaphthalen) is a fluorescent probe that is sensitive to the presence of water molecules in membranes, thereby allowing inference on membrane fluidity. We calculated the generalized polarization (GP) from the fluorescence intensity at 440 and 490 nm emission (Brankatschk et al., 2018; Kaiser et al., 2009) and found that in controls, the GP decreases upon refeeding, suggesting an increase in membrane fluidity. In pex19 mutants, the GP does not change between starved and fed animals, although it is lower than in starved controls (Figure 6F). We hypothesize that in the absence of Pex19, Golgi-peroxisome interaction is impaired, thereby preventing peroxisome-mediated alterations of Golgi membranes required for efficient DCV secretion, such as withdrawal and oxidation of longer chain fatty acids, upon refeeding. In the absence of Pex19 and its direct interactors, such as Pex3 and Pex16, but not proteins involved in matrix protein import, ceramides accumulate in the Golgi, Golgi proteins are missorted into DCVs and membrane fluidity no longer adapts during refeeding, thereby ultimately preventing efficient secretion of insulin-like peptides (Figure 6G).

## Discussion

The secretion of insulin in response to nutritional stimuli such as glucose, amino acids and free fatty acids, is a vital mechanism ensuring the homeostasis of blood glucose levels (Seino et al., 2011). Systemic signals induce insulin secretion from pancreatic β-cells: for example, nutrient uptake in the gut leads to the release of incretin hormones such as Glucagon-like peptide 1 that potentiate insulin secretion compared to direct glucose injection into the blood (Nauck & Meier, 2016). Also the action of insulin itself is systemic: it is secreted into the blood stream, regulating e.g. glucagon release from α-cells in the pancreas and glucose uptake into various organs, e.g. muscles (Tokarz et al., 2018). On the subcellular level, binding of insulin to its receptor in the plasma membrane triggers a signaling cascade including the phosphorylation of phosphoinositides and culminating in the regulation of transcription factors such as FoxO (Lee & Dong, 2017). A crucial subcellular structure in insulin secretion is the Golgi apparatus: it sorts proinsulin into secretory granules, where it is cleaved into mature insulin. Here we present evidence for a cell-intrinsic regulatory mechanism of insulin secretion at the Golgi that reacts to nutrient stimuli but acts independent of other signals from the gut or adipose tissue.

In our study we show that peroxisomes interact with the Golgi apparatus to enable the secretion of insulin-like peptides upon a nutrient stimulus. An interaction between these organelles has been shown in yeast, but so far not in metazoans. We link this interaction to Pex19, a multifunctional protein that is involved in *de novo* peroxisome biogenesis by mediating the budding of vesicles from ER membranes (Agrawal & Subramani, 2016) and in lipid droplet biogenesis (Schrul & Kopito, 2016). Using various lipid biochemical approaches, we demonstrate that medium-and long-chain fatty acids are required in ceramides and that longer fatty acids are removed from DCVs upon refeeding. This mechanism is unfunctional in the absence of Pex19. While we characterized the fatty acid imbalance of pex19 mutants in our previous study (Sellin et al., 2018), here we provide not only several advanced lipidomics and click chemistry analyses, demonstrating a lipid class specific effect of peroxisome metabolism, but also mechanistically untangle a novel function of Pex19 and peroxisomes in neuropeptide secretion. Thereby we provide a new concept for the fundamental cellular process of secretion. Moreover, we demonstrate this in in cells in their native tissue context of a metazoan *in vivo* model.

The secretion of insulin is bi-phasic: a small portion of vesicles is present at the plasma membrane for rapid release, while more secretory granules are shielded from the plasma membrane by actin fibers and require cytoskeleton rearrangement to allow their access to the plasma membrane (Tokarz et al., 2018). While we demonstrate impaired Dilp2 secretion in pex19 mutants following refeeding, it remains possible that the secretion is not completely blocked but rather inefficient and happening at slower kinetics. Upon longer refeeding, Dilp2 might be cleared from pex19 mutant IPC somata eventually. This is supported by our finding that dietary MCFA rescue the secretion of Dilp2 in pex19 mutants, although Pex19 and peroxisomes are not restored under this condition. It is further supported by our finding that the electrophysiological activity of IPCs in pex19 knock-down flies recovers upon refeeding with glucose, suggesting that the neuronal machinery remains functional. It is noteworthy that in the absence of Pex19, neuronal activity of IPCs is uncoupled from insulin-like peptide secretion, while in the presence of peroxisomes, IPC electrical activation has been directly linked to circulating Dilp2 levels (Park et al., 2014).

The mechanism that we describe here is probably not restricted to insulin-like peptide secretion, but rather a general cellular phenomenon. Further studies could focus on other peptidergic neurons in the *Drosophila* brain or other secretory cells, for example in the gut. A further explanation for the impaired Dilp2 secretion and altered DCV properties in pex19 mutants could be a direct association of Pex19 in DCVs. Pex19 is known to function as a chaperone for membrane proteins, and it could potentially interact with proteins on the DCV membrane independently of its role in peroxisome biogenesis (Emmanouilidis et al., 2017; Oh et al., 2025). This hypothesis is supported by the observation that only peroxins involved in membrane protein import (Pex19, Pex3, Pex16), but not those involved in matrix protein import (Pex1, Pex2), affect Dilp2 secretion. The enrichment of GM130 in DCVs from pex19 mutants further suggests potential missorting of membrane proteins that could be directly regulated by Pex19.

A significant finding is the accumulation of phosphatidylinositol (PI) and increased incorporation of C14 fatty acids into PI in pex19 mutants. However, the direct connection between these lipid alterations and impaired Dilp2 secretion remains to be established. PI serves as a precursor for phosphoinositides, which are crucial regulators of membrane trafficking and vesicle fusion. The observed PI accumulation could lead to dysregulation of phosphoinositide levels, particularly PI4P, which has been implicated in Golgi function and vesicle formation (Shinoda et al., 2024). An interesting follow-up project would be to determine phosphoinositide lipidomics in pex19 mutants for more insights into how peroxisomal dysfunction impacts signaling lipids and vesicle trafficking. While our study focuses on Golgi dynamics, the accumulation of PI may also reflect broader changes in ER homeostasis. The ER is the primary site of PI metabolism and ceramide synthesis (Y. J. Kim et al., 2011; Rodriguez-Gallardo et al., 2020).

Ultimately, while we convey some key findings into human HeLa cells, it would be interesting to transfer the role of Pex19 in insulin secretion in pancreatic β-cells. Further studies could employ for example MIN6 cells, a mouse pancreatic beta-cell line, to assess the conservation of the mechanisms reported in our study.

## Material and Methods

### Fly husbandry

Flies were reared on standard cornmeal food (130 g of yarn agar, 248 g of Baker’s yeast, 1223 g of cornmeal, and 1.5 liters of sugar beet syrup in 20 liters of distilled water) and kept in a 25°C incubator with light-dark cycle. The fly stocks obtained from Bloomington Drosophila Stock Center (BDSC) were w^1118^ (no. 6326), dilp2-Gal4 (no. 37516), pex3^2^ (no. 64251), pex2HP^35039^ (no. 21973), pex2^f01899^(no. 18485), Grasp65-GFP (no. 8507), Df(32)6262 (Sellin et al., 2018), UAS-GFP-SKL (no.28882), UAS-Golgi-RFP (no. 30907). The fly stocks obtained from Vienna Drosophila Resource Center (VDRC) were Pex16-RNAi (no. 34296) and Pex1-RNAi (no.27743). UAS mCD8-GFP, UAS PI3K, and UAS PTEN were kindly provided by Reinhard Bauer’s lab and the UAS PLC-delta-PH-GFP stock was kindly provided by Gabriela Edwards Faret. The group of Michael Pankratz (University of Bonn, Germany) kindly provided the stock dilp2,3-Gal4 and UAS-LacZ flies were kindly provided by Carl Thummel (University of Utah, Salt Lake City).

To generate a transgenic construct expressing a peroxisome-targeted mCherry, the coding sequence of mCherry was amplified by PCR from the plasmid pBSKI2_Crest3_insulators-mCherry (Gift from Kathrin Simon, University Bonn) and the peroxisomal targeting signal 1 (PTS1) was cloned to the C-terminus using these primers: fwd: 5’-CAAAATGGTGAGCAAGGGCGAGG-3’; rev: 5’-TTACAACTTCGACTTAGTCTCAGGCGGGTTCTTCTTGTACAGCTCGTCCATGCC-3‘. A second PCR was performed on the purified amplicon (size 745 bp) to generate 5’-overhangs: fwd 5’-GAATTGGGAATTCGTTAACACAA AATGGTGAGCAAGGGCGAGG-3’; rev: 5’-ACCCTCGAGCCGCGGCCGCATTACAACTTCGACTTAGTCTC-3’. The final purified amplicon (size 785 bp) was assembled into a BgIII-linearized and dephosphorylated pUASTattB vector backbone using Gibson assembly (New England Biolabs). Correct integration of insert was verified by test digestion with EcoRI and XbaI. Positive clones were identified and sequenced using these primers: fwd: 5’-GCGCCGGAGTATAAATAGAGG-3’; rev: 5’-TCATCAGTTCCATAGGTTGGAATC-3’.

Two independent pex19 mutant alleles, pex19^ΔF7^ (Bülow et al., 2018) and pex19^ΔB2^, both balanced over CyO with a fluorescent marker (twist-GFP), were crossed to generate transheterozygous Pex19 mutants. Homozygous larvae were selected for absence of the fluorescent marker.

### Starvation and refeeding assay

For the starvation and refeeding assay, w^1118^ control and pex19^-/-^ larvae were reared from the 1^st^ larval stage on apple juice plates (2g Agar Kobe I [Roth], 2.5 g sucrose, 25 mL apple juice, 75 mL water) with additional yeast paste (1 cube [42 g] fresh yeast with 10 mL water) at 25 °C. As early 3^rd^ instar larvae, they were transferred to petri dishes with PBS-soaked filter paper for 2 h. Subsequently, half of the larvae were used immediately for dissection of the tissue and subsequent immunostaining. 10 % sucrose solution in water and a spot of fresh yeast paste 1 h at 25°C were added to the filter paper to provide sugar and protein for refeeding. For MCFA feeding, larvae were reared from the 1^st^ larval stage on apple juice agar plates, and 5 % coconut oil was mixed with the yeast paste. Starvation and refeeding were performed as described above. Larval brains were dissected and immunostaining was performed (see Imaging section).

For starvation and refeeding of adult flies, 3 day old flies were starved for 24 h on filter paper soaked with PBS. Further PBS was added during this time to prevent desiccation. Flies were refed for 4 h with 20 % D-Glucose before dissection.

### Western blot

For the detection of Dilp-2 and GM130, whole larvae homogenates and enriched DCV samples were analyzed. Laemmli buffer (1x) was added, and the lysates were boiled at 70°C for 10 min to destroy all proteases. Electrophoresis was performed at 200V in 1X SDS running buffer. Wet tank blotting was performed to transfer the proteins onto a polyvinylidene difluoride (PVDF) membrane. Blotting was done at 100V for 1 hour on a prior in methanol-activated membrane. The membrane was blocked in 5% BSA in tris-buffered saline with Tween 20 (TBS-T) buffer (1M) for 1 hour followed by incubation with anti-Dilp-2 (1:1,000; gift from M. Pankratz) or anti-GM130 (1:1,000; Abcam Ab30637) antibody in 5%BSA/TBS-T buffer at 4°C overnight. For immunodetection of the antibody, membrane was incubated with horseradish peroxidase (HRP)-coupled secondary antibody (1:10,000; Jackson ImmunoResearch) in 5 % BSA/TBS-T buffer in the dark at room temperature for 2 hours. Detection of antibodies was done with Pierce^TM^ ECL Western Blotting Substrate (Thermo Fisher Scientific) incubated for 1 min and exposed for 20 min. Signal acquisition was performed with Amersham^TM^ Imager 600.

### Electrophysiology of adult flies

All experiments were performed in mated females between 3-6 days post eclosion. Flies were cold anesthetized on ice and then immobilized in a custom-made shim plate fly holder using UV glue (Proformic C1001, VIKO UG, Munich, DE). The proboscis was glued to the thorax to restrict brain movement, and the front legs were excised. The cuticle was then removed in a window above the pars intercerebralis so that the posteriordorsal part of the fly brain was exposed. Trachea and the ocellar ganglion were removed to render the IPCs accessible. The fly was then transferred with the fly holder into a customized, upright fluorescence microscope setup (Olympus BX51WIF, Evident Corporation, Tokyo, JPN). Live images of the fly brain were acquired with a high-resolution camera (SciCam Pro, Scientifica, Uckfield, UK) using an image acquisition software (OCULAR™, Digital Optics Limited, Auckland, NZ).

During the preparation, as well as the experiment, the brain was continuously perfused with carbonated (95% O2 and 5% CO2) extracellular saline containing, 103 mM NaCl, 3 mM KCl, 5 mM N-[Tris(hydroxymethyl)methyl]-2-aminoethanesulfonic acid, 20 mM sucrose, 26 mM NaHCO3, 1 mM NaH2PO4, 1.5 mM CaCl2.2H2O, 4 mM MgCl2.6H2O, and osmolarity adjusted to 273-275 mOsm (modified from (Gouwens & Wilson, 2009). 0.025% Collagenase (w/v in extracellular saline, Sigma-Aldrich #C5138) was gently applied using a thin-walled glass pipette to dissociate the neural sheath above cell bodies before recordings. IPC/DH44^PI^N cell bodies were identified via GFP expression and whole-cell patch clamp recordings were performed using thickwalled patch pipettes (4 – 8 MΩ resistance) containing intracellular saline (40 mM potassium aspartate, 10 mM HEPES, 1 mM EGTA, 4 mM MgATP, 0.5 mM Na3 GTP, 1 mM KCl and 20 μM, 265 mOsm, pH 7.3). Continuous time series recordings of the membrane potential were captured in current clamp mode with an AxoPatch MultiClamp 200B (Molecular Devices, Sunnydale, CA, USA) and corrected for a 13 mV liquid junction potential (Gouwens & Wilson, 2009). All data were recorded with a Digidata 1440A analog-digital converter (Molecular Devices), controlled by the pCLAMP 10 software using a 10 kHz low-pass filter and a 20 kHz sampling rate. Recordings were accepted for analysis if the resting membrane potential was < –48 mV and the spike amplitude > 20 mV. Baseline activity was always recorded in glucose-free saline for 10 min, and the last five min were used for analysis.

### Cell Culture and Immunofluorescence Staining

HeLa cells were maintained in T75cm² flasks using Dulbecco’s Modified Eagle’s Medium (DMEM) high glucose supplemented with 1× penicillin-streptomycin and 10% fetal calf serum (FCS). Cells were cultured at 37°C in a humidified atmosphere containing 5% CO₂. Cells were passaged three times per week at ratios of 1:3 to 1:5 depending on confluence. All experiments were conducted using cells between passages 3 and 10.

For immunofluorescence staining, cells were seeded in 8-well chamber slides (Sarstedt) at a density of 80,000 cells per well. Wells were pre-coated with fibronectin (0.5 µg/µL) for 1 hour at room temperature prior to cell seeding. The staining procedure was performed as follows: Culture medium was aspirated and cells were briefly washed with HBSS containing 10 mM HEPES buffer (HBSS/HEPES). Cells were then incubated with 5 µM BODIPY-C5 (Invitrogen) or 5µM NBD-C6 (Thermo Fisher Scientific) in HBSS/HEPES for 30 minutes at 4°C. After removing the BODIPY solution, fresh HBSS/HEPES buffer was added and cells were incubated at 37°C for 30 minutes. Following a brief wash with HBSS/HEPES buffer, cells were fixed with 4% paraformaldehyde in HBSS/HEPES for 15 minutes at room temperature.

For immunostaining, cells were permeabilized and blocked in HBSS/HEPES containing 0.1% saponin and 10% donkey serum for 1 hour at room temperature. Primary antibodies were PMP70-rabbit (gift from H. Weiher) or GM130-rabbit (Abcam) both at 1:1000 dilution. They were applied in HBSS/HEPES and incubated overnight at 4°C. The following day, cells were washed three times for 5 minutes each with 0.1% saponin in HBSS/HEPES at room temperature. Secondary antibody (anti-rabbit-635) and phalloidin-555 (both at 1:250 dilution) were applied in HBSS/HEPES and incubated for 2 hours in the dark at room temperature. After three 5-minute washes with 0.1% saponin in HBSS/HEPES, nuclei were counterstained with DAPI (1:10,000) in HBSS/HEPES for 5 minutes at room temperature. Following a brief wash with 0.1% saponin in HBSS/HEPES, slides were mounted using Fluoromount (Invitrogen) and stored at 4°C until imaging.

### Fatbody and gut staining

For immunofluorescence staining, fat body and gut tissue from third instar larvae were dissected in cold PBS. The staining procedure was performed as follows: PBS was removed and tissue was incubated with 5 µM BODIPY-C5 (Invitrogen) in PBS for 30 min at 4°C. After removing BODIPY solution, fresh PBS was added and tissue was incubated at 25°C for 30 min. Following a brief wash with PBS, tissue was fixed with 4% paraformaldehyde in PBS for 30 min at room temperature. Afterwards the tissue was permeabilized and blocked with 0.1% saponin and 10% donkey serum in PBS for 30 minutes at RT. F-actin was stained by incubation with Alexa-Fluor-555-phalloidin (Life Technologies; 1:250) in PBS for 2 hours in the dark at RT. After three 15 min washes with 0.1% saponin in PBS, nuclei were counterstained with DAPI (1:10,000) for 5 min at RT. Following a brief wash with 0.1% saponin in PBS, tissue was mounted in Fluoromount (Invitrogen) and stored at 4°C until imaging.

### Whole larvae extraction, DCV purification and lipid extraction for lipidomics analysis

For whole larvae extraction, individual third instar larvae were homogenized in 150 µL LC-MS grade water using a Potter homogenizer. Dense core vesicle (DCV) purification was performed as previously described (Birinci et al., 2020) with modifications. Briefly, third instar larvae were rinsed in PBS to remove any residual food particles. Larvae were then homogenized in 300 µL DCV buffer (300 mM sucrose, 20 mM HEPES pH 7.4) and centrifuged at 1,000 g for 15 min at 4°C. The resulting pellet was resuspended in 300 µL fresh DCV buffer and centrifuged again at 1,000 g for 15 min at 4°C. The supernatant was collected and further centrifuged at 12,000 g for 15 min at 4°C. The final pellet was resuspended in 100 µL DCV buffer for downstream analysis. For lipidomic analysis, lipids were extracted from either whole larvae homogenates or enriched DCVs using a modified protocol based on Thiele et al. (2019). A 50 µL aliquot of either sample was combined with 500 µL extraction mix [5/1 methanol/CHCl3 (LC-MS grade)] + 20 µl OMICS internal standard. Samples were sonicated for 30 min in a bath sonicator and centrifuged for 2 min at 20,000 g. The supernatant was decanted into a fresh tube and 300 µl CHCl3 and 600 µl 1% acetic acid in LC-MS grade water were added. Samples were shaken manually for 10 s and centrifuged at 20,000 g for 5 min. The lower phase was transferred into a fresh tube and dried in a speed-vac (Eppendorf Concentrator Plus) for 15 min at 45°C and redissolved in 500 µl spray buffer [8/5/1 isopropanol/methanol/H2O (all LC-MS grade) + 10 mM ammonium acetate + 0.1% acetic acid (LC-MS grade)] until mass spectrometry analysis.

### Laurdan Assay

Membrane fluidity was assessed using Laurdan-based fluorescence assay adapted for *Drosophila* third instar larvae. Larvae were prepared under experimental conditions of interest (e.g. starved and refed). Prior to assay weight of larvae was detected. Fifteen larvae were selected per biological replicate, washed with distilled water, dried briefly on sterile tissue paper and transferred to Eppendorf cups. Laurdan (6-Dodecanoyl-2-Dimethylaminoaphtalene; stock 5 mM in DMSO) was equilibrated to room temperature. Larvae were homogenized in 300 µL cold DCV buffer (300 mM sucrose, 20 mM HEPES pH 7.4) using a pestle. Homogenates were clarified by centrifugation at 1,000 g for 15 min at 4°C. The resulting supernatant was transferred to fresh Eppendorf cups and further centrifuged at 12,0000 g for 15 min at 4°C to isolate membrane-enriched fractions. The pellet was resuspended in 300 µL fresh DCV buffer containing 3 µL of Laurdan, yielding a final concentration of 50 µM. Samples were incubated at 25°C for 30 min in the dark. Following incorporation, samples were centrifuged at 1,000 g for 15 min and the supernatant was transferred to a new tube. A final spin at 12,000 g for 15 min was performed to pellet the labeled membranes. For fluorescence analysis, the final pellet was resuspended in 100 µL DCV buffer. For experiments requiring subsequent protein analysis (e.g. Western Blot), pellets were alternatively resuspended in 100 µL 1x Complete buffer (Roche). Samples were pipetted into black-walled, clear flat-bottom 96-well microtiter plates in a total volume of 200 µL per well. Fluorescence measurements were carried out using a Tecan plate reader with excitation at 385 nm and emission scan from 400 to 600 nm. The generalized polarization (GP) was calculated: GP = (I440 – I490)/(I440 + I490) with I as the fluorescence intensity (Kaiser et al., 2009).

### Alkyne lipid transfer analysis by mass spectrometry

A previously published protocol (Thiele et al., 2012) to preload Drosophila tissue with alkyne myristate was optimized (Carrera et al., 2024). Click reaction and mass spectrometry analysis were carried out as previously described (Thiele et al., 2019; Wunderling et al., 2023). Briefly, whole larval tissue from control and Pex19 mutant larvae was dissected in triplicate in hemolymph-like buffer (HL3A; 115 mM sucrose, 70 mM NaCl, 20 mM MgCl2, 10 mM NaHCO3, 5 mM KCl, 5 mM HEPES, 5 mM trehalose, pH 7.2; (Paradis et al., 2022). Tissue was preloaded with 50 µM alkyne myristate (FA 14:0;Y) in HL3A when incubated on a nutator for 60 min. Tissue was washed three times in HL3A. Lipids were extracted from tissue as described above but additional internal standards for alkyne labeled metabolites were included (Thiele et al., 2019). After drying the sample, lipids were resolved in 10 µl CHCl3 and sonicated for 5 min. Then 40 µl of C171 click mix [prepared by mixing 10 µl of 100 mM C171 in 50% methanol (stored as aliquots at −80°C) with 200 µl 5 mM Cu(I)AcCN4BF4 in AcCN and 800 µl ethanol] were added and the samples were incubated at 40°C overnight (16 h). After this, 200 µl CHCl3 and 200 µl water were added and the samples were centrifuged at 20,000 g for 5 min. The upper phase was discarded and samples were dried in a speed-vac for 15 min at 45°C. Samples were resolved in 500 µl spray buffer and analyzed by mass spectrometry.

### Imaging

Antibodies used in this study were α-GFP-mouse (Santa Cruz Biotechnology), α-DsRed-rabbit (Takara), α-dilp2-guinea pig (Gift from M. Pankratz), α-dilp3-rabbit (kindly provided by Hannah Rindsfüßer), α-GM130-rabbit (Abcam), α-PMP70-rabbit (Gift from H. Weiher). For immunohistochemistry, brains from larvae were dissected in PBS and fixed for 1 hour in 0.5% PBS-Tween 20 and 4% formaldehyde. Tissue was washed with 0.5% PBS-Tween 20 and blocked with donkey serum before incubation with the primary antibody (overnight at 4°C). Tissue was washed in 0.1% PBS-Tween 20 before incubation with the secondary antibody at room temperature in the dark for 1 hour. Secondary antibodies coupled to Alexa dyes were from Biozol. Tissue was washed, incubated for 5 min with DAPI and mounted in Fluoromount (Thermo Fisher Scientific). Secondary antibodies were donkey-α-mouse-Alexa488 and donkey-α-rabbit-Cy3. For imaging, we used a Zeiss LSM 880 with Airyscan detector and a 20x air lens, 40x oil lens and 63x oil lens (all Plan-Apochromat, Zeiss). For quantification of fluorescence intensity, we used ImageJ on images taken with the same laser intensity and gain. Fluorescence intensity was determined as the background-corrected intensity of individual IPC somata. A minimum of 3 brains was analyzed per condition.

### Imaris analysis of Peroxisome-Golgi interaction

Image analysis was carried out using Imaris version 9.6.1, specifically the modules *ImarisCell*, *Spots*, and *Surfaces*, to investigate peroxisome-Golgi interaction dynamics and organelle morphology. Prior to analysis, sample conditions (starved vs. refed) were blinded to eliminate bias. A representative file was selected to establish a batch-processing pipeline consisting of the following set of objects: First, the *Cell* function of *ImarisCell* was utilized to identify the somata of the IPCs within the cluster. In order to exclude staining artifacts outside the IPC cluster and to remove signal originating from axonal compartments, signal channels were masked using a *Surface* object generated from the segmented somata. All subsequent analysis steps were performed on these masked channels. For the Golgi apparatus, *Surfaces* were created based on the Golgi-RFP signal using the following parameters: *smoothing* enabled, *surface grain size* set to 0.181 µm, *background elimination* enabled, and the *diameter of the largest sphere* set to 0.679 µm. For the peroxisomal GFP-SKL signal, *Spots* were generated with an *estimated diameter* of 0.500 µm*, background subtraction* enabled, and the *region growing* feature activated. The *region growing* function was employed to generate *Spots* of varying sizes based on the underlying signal. The *region growing* function was based on *local contrast* and the diameter on *region border*. The batch job was executed on all files, and subsequent modifications that could not be performed in batch mode were carried out on each file individually. The display of all channels was automatically adjusted, and additional adjustments were made to accurately represent the signal. These adjustments included modifying the threshold value of the *Quality Filter* in the subsection Filters of the *Creation Wizard* for *Spots* and *Surfaces*. Cells that exhibited insufficient signal quality or expressed only one of the two markers were excluded from further analysis. Exported parameters included *Spots-Total Number of Spots, Spots-Volume, Spots-Average Distance To 3 Nearest Neighbour, Spots-Shortest Distance to Surfaces* and *Surface-Sphericity, Surfaces-Total Number of Surfaces, Surfaces-Volume, Surfaces-Shortest Distance to Surfaces*. Calculation of the arithmetic mean, the sum of the volumes of objects, the percentage of colocalizing objects, and the normalization to the manually determined cell count was done in Microsoft Excel. Colocalization was defined as a shortest distance of less than or equal to 0 µm between a peroxisomal *Spot* and a Golgi *Surface*.

### Statistics

Bar charts represent mean and standard deviation. Boxes in box plots represent the interquartile range and median, whiskers represent minimum and maximum. Green dots in box plots and bar charts represent single data points. We used Graph Pad Prism for graphs and statistical analyses. Two-sided Student’s t-test was applied for normally distributed data in single comparisons, assuming heteroschedasticy. One-way ANOVA with Tukey-Kramer post-test was used for multiple comparisons. The Kolmogorov-Smirnow test was applied to test normality, and Bartlett’s method was used to test for equal standard deviations within groups. Asterisks represent * = p < 0.05, ** = p < 0.01, *** = p < 0.001. A minimum of 3 biological replicates was used for each analysis.

### Visualization

We used biorender for illustrations of protein and organelle interaction.

## Author contribution

Conceptualization: MHB; Methodology: MHB, JMA, KW, LK, CT, TRB; Investigation: MHB, MAK, NK, DPDR, HS, KW, TRB, MY; Writing – Original Draft: MHB, NK, DPDR; Resources – MHB; Writing – Review & Editing: MHB, NK, DPDR, MAK; Visualization: MHB, MAK, NK, DPDR; Funding Acquisition: MHB; Supervision: MHB, JMA, LK.

## Acknowledgments

We thank Markus Islinger, Julia Sellin, Hirak Das, Johannes Broichhagen and Lutz Schmitt for valuable discussions. We thank Damian Demarest, Andreas Schoofs, Tania Groß and Michael Pankratz for sharing the dilp2 antibody and fly lines, Bianca Schrul for sharing PEX19 mutant HeLa cells and Hans Weiher for the PMP70 antibody. We thank Naoki Kato, Julia Kalinowski, Yllka Ahmeti, Santiago Maya and Yerin Jung for help with experiments. We thank Reinhard Bauer, Franka Eckardt, Elvira Weber, Hug Aubin and Artur Lichtenberg for their support and all members of the CURE3D research lab of the University Hospital Düsseldorf for valuable discussions. We thank the AdLight Imaging facility of the Medical Faculty of the University of Düsseldorf. MHB received funding from the German Research Foundation (DFG, project numbers 417982926 and 535112684), from the University of Bonn (TRA Matter and TRA Life & Health INNOVATION grant) and from the Medical Faculty of the University of Düsseldorf (Integration grant).

## Supplement

**Supplemental figure S1:**
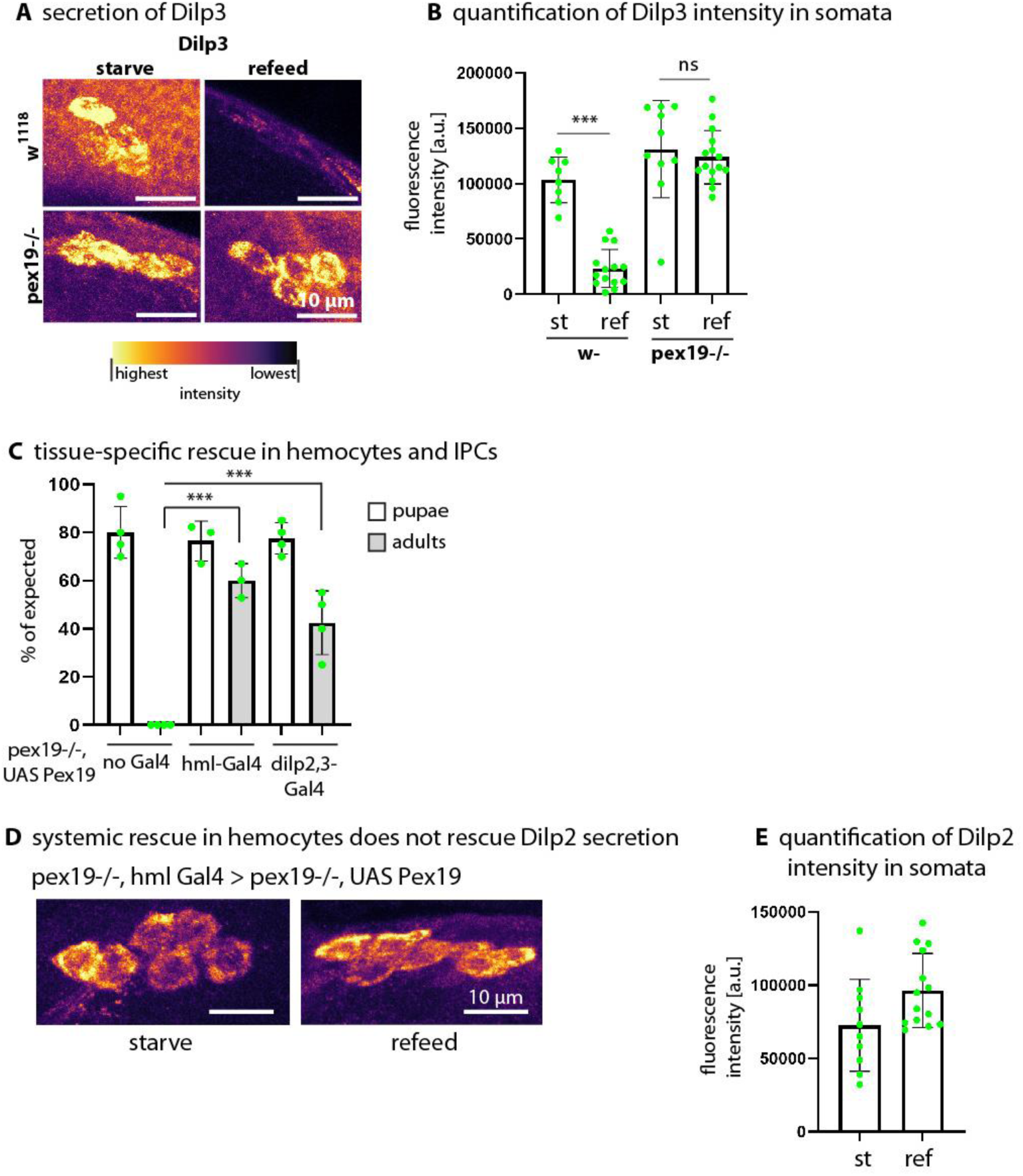
(A) Secretion of Dilp3 in insulin-producing cells (IPCs). Representative images of larval IPCs in wild-type (w^1118^) and pex19 mutant (pex19^ΔF7/ΔB2^), comparing starved and refed states. Heatmap indicates Dilp3 fluorescence intensity (highest to lowest). (B) Quantification of Dilp3 fluorescence intensity in IPCs across conditions in wild-type and pex19 mutants. (C) Tissue-specific rescue of Pex19 in hemocytes and IPCs. Percentage of expected pupae and adult flies in pex19^−^/^−^, UAS-Pex19 crosses with different Gal4 drivers: no Gal4 (genotype: w-; pex19-/-, UAS Pex19; +), hml-Gal4 (genotype: w-; pex19-/-, hml-Gal4/UAS Pex19; +), and dilp2,3-Gal4 (genotype: w-; pex19-/-, UAS Pex19; dilp2,3 Gal4). n ≥ 3 in groups of 25 animals. (D) Systemic rescue in hemocytes does not restore Dilp2 secretion. Representative images of IPCs in pex19-/-, hml-Gal4/UAS Pex19 larvae under starved and refed conditions. (E) Quantification of Dilp2 fluorescence intensity in IPCs across conditions. Scale bars as indicated. Asterisks represent *** p < 0.001, ns: not significant.

**Supplemental Figure S2:**
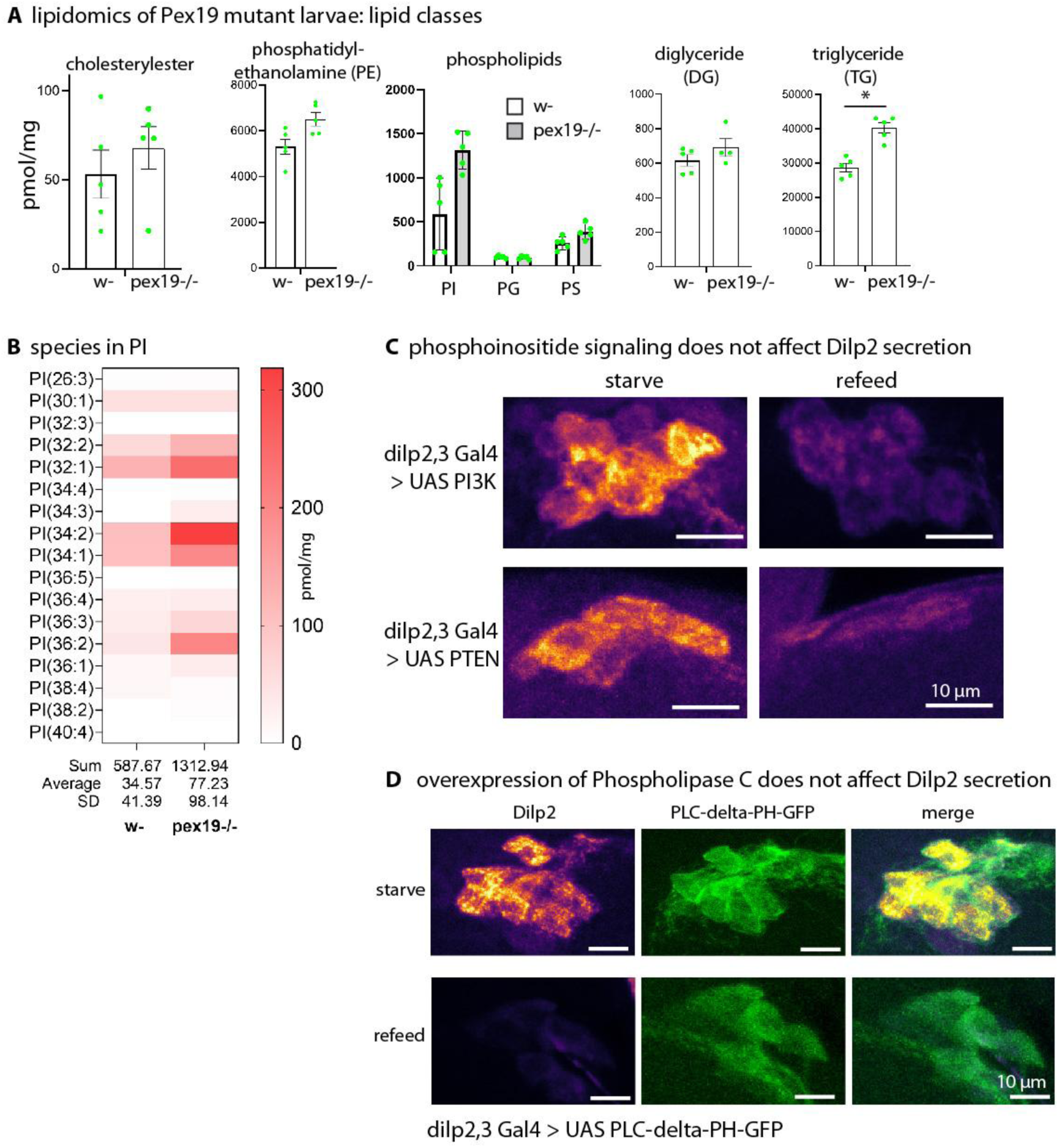
(A) Lipidomics of w- and pex19 mutant larvae separated by different lipid classes: cholesterylester, PE, other phospholipids and the storage lipids DG and TG. (B) Distribution of lipid species in PI. (C) Confocal images of larval IPCs stained against Dilp2. Overexpression of PI3K and PTEN in IPCs does not affect Dilp2 secretion. Genotypes are +; dilp2,3 Gal4/UAS PI3K and +; dilp2,3 Gal4/UAS PTEN (D) Expression of PLC-δ- PH-GFP does not affect dilp2 secretion. Genotype: dilp2,3 Gal4/UAS PLC-δ-PH-GFP. Scale bars as indicated. Asterisks represent * p < 0.05.

**Supplemental Figure S3:**
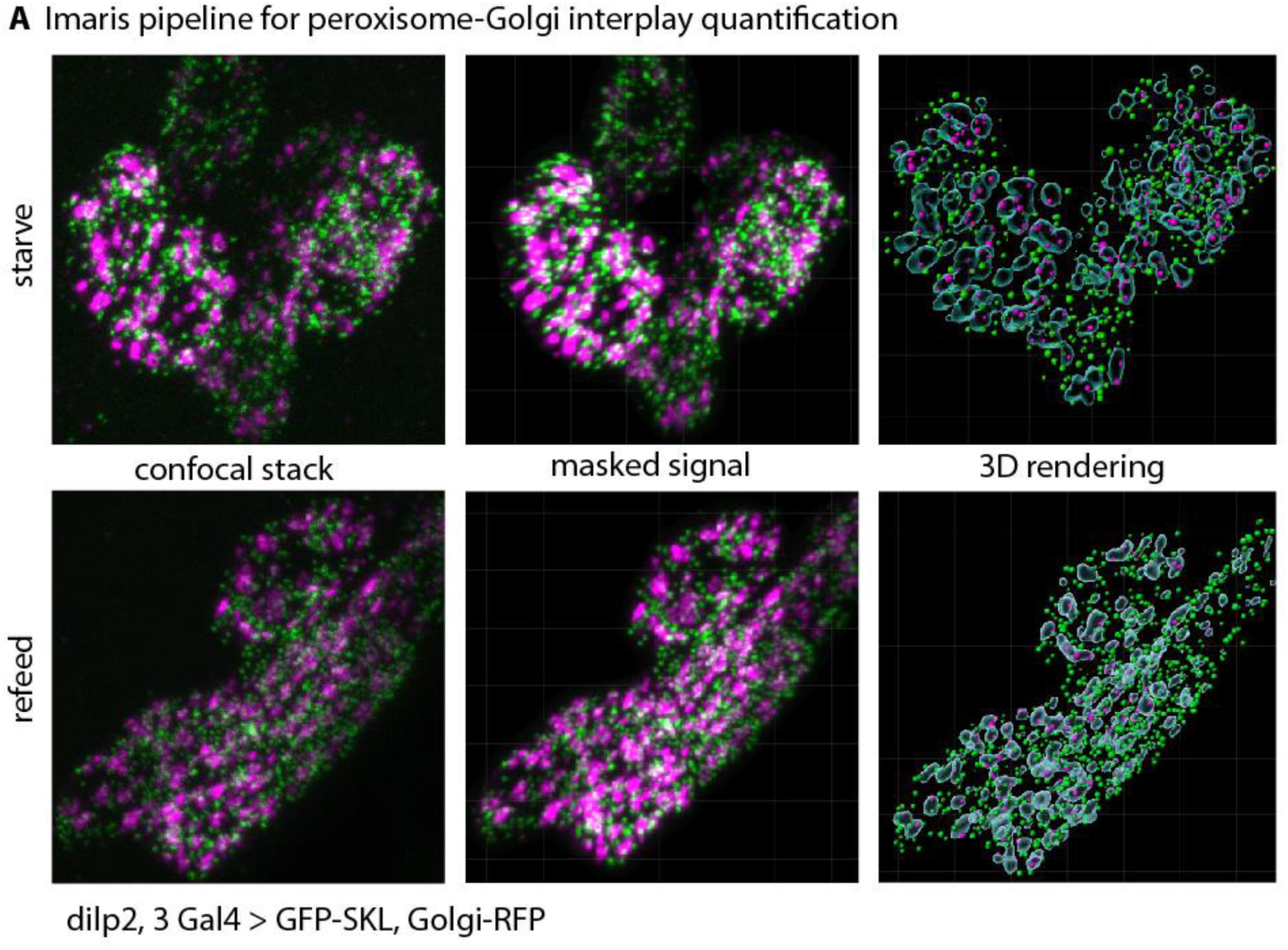
(A) Imaris pipeline to create a masked signal from confocal z-stacks and subsequent 3D rendering.

**Supplemental Figure 4:**
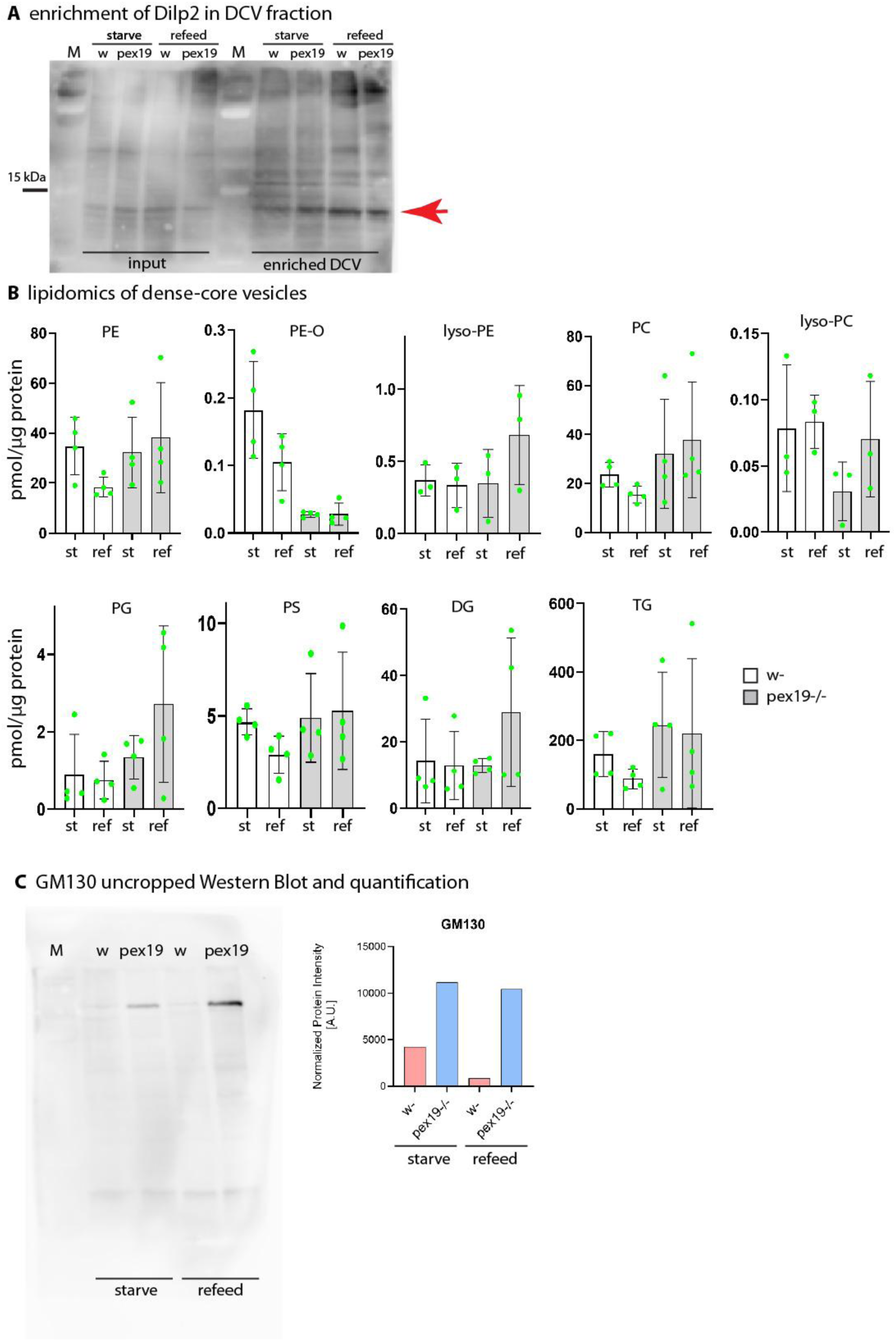
(A) Western Blot showing enrichment of Dilp2 compared to input (larval extracts) after the DCV enrichment protocol. 17 μg protein were loaded to each lane. Dilp2 has a size of ∼ 10 kDa (indicated by red arrow). (B) Lipidomics analysis of enriched DCVs from control and pex19 mutants upon starvation and refeeding. (C) Uncropped Western Blot of GM130 in DCV fractions and quantification, normalized to protein input.

## References

Agliarulo, I., & Parashuraman, S. (2022). Golgi Apparatus Regulates Plasma Membrane Composition and Function. Cells, 11(3), 368. 10.3390/cells11030368

Agrawal, G., & Subramani, S. (2016). De novo peroxisome biogenesis: Evolving concepts and conundrums. Biochimica et Biophysica Acta (BBA) - Molecular Cell Research, 1863(5), 892–901. 10.1016/j.bbamcr.2015.09.014

Baboota, R. K., Shinde, A. B., Lemaire, K., Fransen, M., Vinckier, S., Van Veldhoven, P. P., Schuit, F., & Baes, M. (2019). Functional peroxisomes are required for β-cell integrity in mice. Molecular Metabolism, 22, 71–83. 10.1016/j.molmet.2019.02.001

Bader, R., Sarraf-Zadeh, L., Peters, M., Moderau, N., Stocker, H., Köhler, K., Pankratz, M. J., & Hafen, E. (2013). The IGFBP7 homolog Imp-L2 promotes insulin signaling in distinct neurons of the *Drosophila* brain. Journal of Cell Science. 10.1242/jcs.120261

Bai, H., Kang, P., & Tatar, M. (2012). Drosophila insulin-like peptide-6 (*dilp6*) expression from fat body extends lifespan and represses secretion of Drosophila insulin-like peptide-2 from the brain. Aging Cell, 11(6), 978–985. 10.1111/acel.12000

Bao, X., Koorengevel, M. C., Groot Koerkamp, M. J. A., Homavar, A., Weijn, A., Crielaard, S., Renne, M. F., Lorent, J. H., Geerts, W. J., Surma, M. A., Mari, M., Holstege, F. C. P., Klose, C., & de Kroon, A. I. P. M. (2021). Shortening of membrane lipid acyl chains compensates for phosphatidylcholine deficiency in choline-auxotroph yeast. The EMBO Journal, 40(20). 10.15252/embj.2021107966

Barry, W. E., & Thummel, C. S. (2016). The Drosophila HNF4 nuclear receptor promotes glucose-stimulated insulin secretion and mitochondrial function in adults. ELife, 5. 10.7554/eLife.11183

Birinci, Y., Preobraschenski, J., Ganzella, M., Jahn, R., & Park, Y. (2020). Isolation of large dense-core vesicles from bovine adrenal medulla for functional studies. Scientific Reports, 10(1), 7540. 10.1038/s41598-020-64486-3

Bisen, R. S., Iqbal, F. M., Cascino-Milani, F., Bockemühl, T., & Ache, J. M. (2025). Nutritional state-dependent modulation of insulin-producing cells in Drosophila. ELife, 13. 10.7554/eLife.98514

Brankatschk, M., Gutmann, T., Knittelfelder, O., Palladini, A., Prince, E., Grzybek, M., Brankatschk, B., Shevchenko, A., Coskun, Ü., & Eaton, S. (2018). A Temperature-Dependent Switch in Feeding Preference Improves Drosophila Development and Survival in the Cold. Developmental Cell, 46(6), 781–793.e4. 10.1016/j.devcel.2018.05.028

Brogiolo, W., Stocker, H., Ikeya, T., Rintelen, F., Fernandez, R., & Hafen, E. (2001). An evolutionarily conserved function of the Drosophila insulin receptor and insulin-like peptides in growth control. Current Biology, 11(4), 213–221. 10.1016/S0960-9822(01)00068-9

Bülow, M. H., & Broichhagen, J. (2025). Chemical probes for imaging cellular compartmentalization. Trends in Biochemical Sciences, 50(2), 171–172. 10.1016/j.tibs.2024.12.005

Bülow, M. H., Wingen, C., Senyilmaz, D., Gosejacob, D., Sociale, M., Bauer, R., Schulze, H., Sandhoff, K., Teleman, A. A., Hoch, M., & Sellina, J. (2018). Unbalanced lipolysis results in lipotoxicity and mitochondrial damage in peroxisome-deficient Pex19 mutants. Molecular Biology of the Cell, 29(4). 10.1091/mbc.E17-08-0535

Carrera, P., Odenthal, J., Risse, K. S., Jung, Y., Kuerschner, L., & Bülow, M. H. (2024). The CD36 scavenger receptor Bez regulates lipid redistribution from fat body to ovaries in *Drosophila*. Development, 151(9). 10.1242/dev.202551

Castro, I. G., Shortill, S. P., Dziurdzik, S. K., Cadou, A., Ganesan, S., Valenti, R., David, Y., Davey, M., Mattes, C., Thomas, F. B., Avraham, R. E., Meyer, H., Fadel, A., Fenech, E. J., Ernst, R., Zaremberg, V., Levine, T. P., Stefan, C., Conibear, E., & Schuldiner, M. (2022). Systematic analysis of membrane contact sites in Saccharomyces cerevisiae uncovers modulators of cellular lipid distribution. ELife, 11. 10.7554/eLife.74602

Chitkara, S., & Atilla-Gokcumen, G. E. (2025). Decoding ceramide function: how localization shapes cellular fate and how to study it. Trends in Biochemical Sciences, 50(4), 356–367. 10.1016/j.tibs.2025.01.007

Emmanouilidis, L., Schütz, U., Tripsianes, K., Madl, T., Radke, J., Rucktäschel, R., Wilmanns, M., Schliebs, W., Erdmann, R., & Sattler, M. (2017). Allosteric modulation of peroxisomal membrane protein recognition by farnesylation of the peroxisomal import receptor PEX19. Nature Communications, 8(1), 14635. 10.1038/ncomms14635

Faust, J. E., Verma, A., Peng, C., & McNew, J. A. (2012). An Inventory of Peroxisomal Proteins and Pathways in *Drosophila melanogaster*. Traffic, 13(10), 1378–1392. 10.1111/j.1600-0854.2012.01393.x

Gouwens, N. W., & Wilson, R. I. (2009). Signal Propagation in *Drosophila* Central Neurons. The Journal of Neuroscience, 29(19), 6239–6249. 10.1523/JNEUROSCI.0764-09.2009

Hänschke, L., Heier, C., Maya Palacios, S. J., Özek, H. E., Thiele, C., Bauer, R., Kühnlein, R. P., & Bülow, M. H. (2022). Drosophila Lipase 3 Mediates the Metabolic Response to Starvation and Aging. Frontiers in Aging, 3. 10.3389/fragi.2022.800153

Ho, K.-H., Jayathilake, A., Yagan, M., Nour, A., Osipovich, A. B., Magnuson, M. A., Gu, G., & Kaverina, I. (2023). CAMSAP2 localizes to the Golgi in islet β-cells and facilitates Golgi-ER trafficking. IScience, 26(2), 105938. 10.1016/j.isci.2023.105938

Honsho, M., Asaoku, S., Fukumoto, K., & Fujiki, Y. (2013). Topogenesis and Homeostasis of Fatty Acyl-CoA Reductase 1. Journal of Biological Chemistry, 288(48), 34588–34598. 10.1074/jbc.M113.498345

Huttlin, E. L., Bruckner, R. J., Navarrete-Perea, J., Cannon, J. R., Baltier, K., Gebreab, F., Gygi, M. P., Thornock, A., Zarraga, G., Tam, S., Szpyt, J., Gassaway, B. M., Panov, A., Parzen, H., Fu, S., Golbazi, A., Maenpaa, E., Stricker, K., Guha Thakurta, S., … Gygi, S. P. (2021). Dual proteome-scale networks reveal cell-specific remodeling of the human interactome. Cell, 184(11), 3022–3040.e28. 10.1016/j.cell.2021.04.011

Huttlin, E. L., Ting, L., Bruckner, R. J., Gebreab, F., Gygi, M. P., Szpyt, J., Tam, S., Zarraga, G., Colby, G., Baltier, K., Dong, R., Guarani, V., Vaites, L. P., Ordureau, A., Rad, R., Erickson, B. K., Wühr, M., Chick, J., Zhai, B., … Gygi, S. P. (2015). The BioPlex Network: A Systematic Exploration of the Human Interactome. Cell, 162(2), 425–440. 10.1016/j.cell.2015.06.043

Iwamoto, T., Shimizu, S., Tajima-Sakurai, H., Yamaguchi, H., Nishida, Y., Arakawa, S., & Watada, H. (2023). Inhibition of Insulin Secretion Induces Golgi Morphological Changes. Juntendo Medical Journal, 69(1), JMJ22-0040-OA. 10.14789/jmj.JMJ22-0040-OA

Jaspers, M. H. J., Pflanz, R., Riedel, D., Kawelke, S., Feussner, I., & Schuh, R. (2014). The fatty acyl-CoA reductase Waterproof mediates airway clearance in Drosophila. Developmental Biology, 385(1), 23–31. 10.1016/j.ydbio.2013.10.022

Kaiser, H.-J., Lingwood, D., Levental, I., Sampaio, J. L., Kalvodova, L., Rajendran, L., & Simons, K. (2009). Order of lipid phases in model and plasma membranes. Proceedings of the National Academy of Sciences, 106(39), 16645–16650. 10.1073/pnas.0908987106

Kim, J., & Neufeld, T. P. (2015). Dietary sugar promotes systemic TOR activation in Drosophila through AKH-dependent selective secretion of Dilp3. Nature Communications, 6(1), 6846. 10.1038/ncomms7846

Kim, Y. J., Guzman-Hernandez, M. L., & Balla, T. (2011). A Highly Dynamic ER-Derived Phosphatidylinositol-Synthesizing Organelle Supplies Phosphoinositides to Cellular Membranes. Developmental Cell, 21(5), 813–824. 10.1016/j.devcel.2011.09.005

Kumar, R., Islinger, M., Worthy, H., Carmichael, R., & Schrader, M. (2024). The peroxisome: an update on mysteries 3.0. Histochemistry and Cell Biology, 161(2), 99–132. 10.1007/s00418-023-02259-5

Lee, R. G., Rudler, D. L., Raven, S. A., Peng, L., Chopin, A., Moh, E. S. X., McCubbin, T., Siira, S. J., Fagan, S. V., DeBono, N. J., Stentenbach, M., Browne, J., Rackham, F. F., Li, J., Simpson, K. J., Marcellin, E., Packer, N. H., Reid, G. E., Padman, B. S., … Filipovska, A. (2024). Quantitative subcellular reconstruction reveals a lipid mediated inter-organelle biogenesis network. Nature Cell Biology, 26(1), 57–71. 10.1038/s41556-023-01297-4

Lee, S., & Dong, H. H. (2017). FoxO integration of insulin signaling with glucose and lipid metabolism. Journal of Endocrinology, 233(2), R67–R79. 10.1530/JOE-17-0002

Liessem, S., Held, M., Bisen, R. S., Haberkern, H., Lacin, H., Bockemühl, T., & Ache, J. M. (2023). Behavioral state-dependent modulation of insulin-producing cells in Drosophila. Current Biology, 33(3), 449–463.e5. 10.1016/j.cub.2022.12.005

Magnan, C., & Le Stunff, H. (2021). Role of hypothalamic de novo ceramides synthesis in obesity and associated metabolic disorders. Molecular Metabolism, 53, 101298. 10.1016/j.molmet.2021.101298

Maruyama, J., & Kitamoto, K. (2013). Expanding functional repertoires of fungal peroxisomes: contribution to growth and survival processes. Frontiers in Physiology, 4. 10.3389/fphys.2013.00177

Nauck, M. A., & Meier, J. J. (2016). The incretin effect in healthy individuals and those with type 2 diabetes: physiology, pathophysiology, and response to therapeutic interventions. The Lancet Diabetes & Endocrinology, 4(6), 525–536. 10.1016/S2213-8587(15)00482-9

Oh, J., Catherine, C., Kim, E. S., Min, K. W., Jeong, H. C., Kim, H., Kim, M., Ahn, S. H., Lukianenko, N., Jo, M. G., Bak, H. S., Lim, S., Kim, Y. K., Kim, H. M., Lee, S. B., & Cho, H. (2025). Engineering a membrane protein chaperone to ameliorate the proteotoxicity of mutant huntingtin. Nature Communications, 16(1), 737. 10.1038/s41467-025-56030-6

Paradis, M., Kucharowski, N., Edwards Faret, G., Maya Palacios, S. J., Meyer, C., Stümpges, B., Jamitzky, I., Kalinowski, J., Thiele, C., Bauer, R., Paululat, A., Sellin, J., & Bülow, M. H. (2022). The ER protein Creld regulates ER-mitochondria contact dynamics and respiratory complex 1 activity. Science Advances, 8(29). 10.1126/sciadv.abo0155

Park, S., Alfa, R. W., Topper, S. M., Kim, G. E. S., Kockel, L., & Kim, S. K. (2014). A Genetic Strategy to Measure Circulating Drosophila Insulin Reveals Genes Regulating Insulin Production and Secretion. PLoS Genetics, 10(8), e1004555. 10.1371/journal.pgen.1004555

Rajan, A., & Perrimon, N. (2012). Drosophila Cytokine Unpaired 2 Regulates Physiological Homeostasis by Remotely Controlling Insulin Secretion. Cell, 151(1). 10.1016/j.cell.2012.08.019

Reumann, S., & Bartel, B. (2016). Plant peroxisomes: recent discoveries in functional complexity, organelle homeostasis, and morphological dynamics. Current Opinion in Plant Biology, 34, 17–26. 10.1016/j.pbi.2016.07.008

Rodriguez-Gallardo, S., Kurokawa, K., Sabido-Bozo, S., Cortes-Gomez, A., Ikeda, A., Zoni, V., Aguilera-Romero, A., Perez-Linero, A. M., Lopez, S., Waga, M., Araki, M., Nakano, M., Riezman, H., Funato, K., Vanni, S., Nakano, A., & Muñiz, M. (2020). Ceramide chain length–dependent protein sorting into selective endoplasmic reticulum exit sites. Science Advances, 6(50). 10.1126/sciadv.aba8237

Sano, H., Nakamura, A., Texada, M. J., Truman, J. W., Ishimoto, H., Kamikouchi, A., Nibu, Y., Kume, K., Ida, T., & Kojima, M. (2015). The Nutrient-Responsive Hormone CCHamide-2 Controls Growth by Regulating Insulin-like Peptides in the Brain of Drosophila melanogaster. PLOS Genetics, 11(5), e1005209. 10.1371/journal.pgen.1005209

Sarraf-Zadeh, L., Christen, S., Sauer, U., Cognigni, P., Miguel-Aliaga, I., Stocker, H., Köhler, K., & Hafen, E. (2013). Local requirement of the Drosophila insulin binding protein imp-L2 in coordinating developmental progression with nutritional conditions. Developmental Biology, 381(1), 97–106. 10.1016/j.ydbio.2013.06.008

Schrul, B., & Kopito, R. R. (2016). Peroxin-dependent targeting of a lipid-droplet-destined membrane protein to ER subdomains. Nature Cell Biology, 18(7), 740–751. 10.1038/ncb3373

Seino, S., Shibasaki, T., & Minami, K. (2011). Dynamics of insulin secretion and the clinical implications for obesity and diabetes. Journal of Clinical Investigation, 121(6), 2118–2125. 10.1172/JCI45680

Sellin, J., Wingen, C., Gosejacob, D., Senyilmaz, D., Hänschke, L., Büttner, S., Meyer, K., Bano, D., Nicotera, P., Teleman, A. A., & Bülow, M. H. (2018). Dietary rescue of lipotoxicity-induced mitochondrial damage in Peroxin19 mutants. PLoS Biology, 16(6). 10.1371/journal.pbio.2004893

Semaniuk, U., Piskovatska, V., Strilbytska, O., Strutynska, T., Burdyliuk, N., Vaiserman, A., Bubalo, V., Storey, K. B., & Lushchak, O. (2021). *Drosophila* insulin-like peptides: from expression to functions – a review. Entomologia Experimentalis et Applicata, 169(2), 195–208. 10.1111/eea.12981

Shinoda, S., Sakai, Y., Matsui, T., Uematsu, M., Koyama-Honda, I., Sakamaki, J., Yamamoto, H., & Mizushima, N. (2024). Syntaxin 17 recruitment to mature autophagosomes is temporally regulated by PI4P accumulation. 10.7554/eLife.92189.2

Thiele, C., Papan, C., Hoelper, D., Kusserow, K., Gaebler, A., Schoene, M., Piotrowitz, K., Lohmann, D., Spandl, J., Stevanovic, A., Shevchenko, A., & Kuerschner, L. (2012). Tracing Fatty Acid Metabolism by Click Chemistry. ACS Chemical Biology, 7(12), 2004–2011. 10.1021/cb300414v

Thiele, C., Wunderling, K., & Leyendecker, P. (2019). Multiplexed and single cell tracing of lipid metabolism. Nature Methods, 16(11), 1123–1130. 10.1038/s41592-019-0593-6

Tokarz, V. L., MacDonald, P. E., & Klip, A. (2018). The cell biology of systemic insulin function. Journal of Cell Biology, 217(7). 10.1083/jcb.201802095

van Galen, J., Campelo, F., Martínez-Alonso, E., Scarpa, M., Martínez-Menárguez, J. Á., & Malhotra, V. (2014). Sphingomyelin homeostasis is required to form functional enzymatic domains at the trans-Golgi network. Journal of Cell Biology, 206(5), 609–618. 10.1083/jcb.201405009

Wangler, M. F., Chao, Y.-H., Roth, M., Welti, R., & McNew, J. A. (2024). Drosophila Models Uncover Substrate Channeling Effects on Phospholipids and Sphingolipids in Peroxisomal Biogenesis Disorders. 10.1101/2024.04.26.591192

Wunderling, K., Zurkovic, J., Zink, F., Kuerschner, L., & Thiele, C. (2023). Triglyceride cycling enables modification of stored fatty acids. Nature Metabolism, 5(4), 699–709. 10.1038/s42255-023-00769-z

Yang, J., Guo, F., Chin, H. S., Chen, G. Bin, Ang, C. H., Lin, Q., Hong, W., & Fu, N. Y. (2023). Sequential genome-wide CRISPR-Cas9 screens identify genes regulating cell-surface expression of tetraspanins. Cell Reports, 42(2), 112065. 10.1016/j.celrep.2023.112065

